# Widespread impact of nucleosome remodelers on transcription at cis-regulatory elements

**DOI:** 10.1101/2024.04.12.589208

**Authors:** Benjamin J. Patty, Sarah J. Hainer

## Abstract

Nucleosome remodeling complexes and other regulatory factors work in concert to build a chromatin environment that directs the expression of a distinct set of genes in each cell using cis-regulatory elements (CREs), such as promoters and enhancers, that drive transcription of both mRNAs and CRE-associated non-coding RNAs (ncRNAs). Two classes of CRE-associated ncRNAs include upstream antisense RNAs (uaRNAs), which are transcribed divergently from a shared mRNA promoter, and enhancer RNAs (eRNAs), which are transcribed bidirectionally from active enhancers. The complicated network of CRE regulation by nucleosome remodelers remains only partially explored, with a focus on a select, limited number of remodelers. We endeavored to elucidate a remodeler-based regulatory network governing CRE-associated transcription (mRNA, eRNA, and uaRNA) in murine embryonic stem (ES) cells to test the hypothesis that many SNF2-family nucleosome remodelers collaborate to regulate the coding and non-coding transcriptome via alteration of underlying nucleosome architecture. Using depletion followed by transient transcriptome sequencing (TT-seq), we identified thousands of misregulated mRNAs and CRE-associated ncRNAs across the remodelers examined, identifying novel contributions by understudied remodelers in the regulation of coding and non-coding transcription. Our findings suggest that mRNA and eRNA transcription are coordinately co-regulated, while mRNA and uaRNAs sharing a common promoter are independently regulated. Subsequent mechanistic studies suggest that while remodelers SRCAP and CHD8 modulate transcription through classical mechanisms such as transcription factors and histone variants, a broad set of remodelers including SMARCAL1 indirectly contribute to transcriptional regulation through maintenance of genomic stability and proper Integrator complex localization. This study systematically examines the contribution of SNF2-remodelers to the CRE-associated transcriptome, identifying at least two classes for remodeler action.

## Introduction

Nucleosome remodeling complexes (remodelers) serve critical roles in DNA-templated processes, including transcription, replication, and DNA repair^1,2^. Remodelers are highly diversified in eukaryotic systems, with at least 32 SNF2-like ATPase proteins representing four subfamilies (SWI/SNF, INO80, ISWI, and CHD) based on the presence of subfamily-specific protein domains within the catalytic subunit^1,2^. Additional SNF2-like ATPase remodelers lacking these defining domains are currently sorted into a less well-defined outgroup^1,2^. Remodelers utilize ATP hydrolysis to translocate DNA, resulting in nucleosome mobilization through various mechanisms which can facilitate or inhibit DNA-templated activities^1,3^.

Several remodelers have been shown to impact protein-coding mRNA expression^1,2^. In murine embryonic stem (ES) cells, esBAF, NuRD, and Tip60-p400 have been characterized as major regulators of mRNA expression that cooperate with and antagonize one another to maintain the pluripotent state of the cell^4–6^. Additional remodelers, including SNF2H, INO80, CHD1, CHD2, CHD8, and SMARCAD1, play smaller yet still important roles in mRNA regulation in ES cells that contribute to pluripotency and other processes^7–15^. However, the contributions of most remodelers to non-protein coding RNA expression, including major transcriptional regulators such as the NuRD and Tip60-p400 complexes, remain unexplored.

Non-protein-coding transcription gives rise to non-coding RNAs (ncRNAs): diverse RNA species, some having well-established roles in transcriptional and translational regulation^16^. Examples of functional ncRNAs include ribosomal RNAs (rRNAs), transfer RNAs (tRNAs), and microRNAs (miRNAs), among others. The function(s) of non-coding transcription originating from cis-regulatory elements (CREs), including promoters (upstream-antisense RNAs, uaRNAs), and enhancers (enhancer RNAs, eRNAs), remains less well defined^16^. Cells must activate networks of enhancers and promoters to drive appropriate mRNA expression in response to signaling cues; in parallel, active enhancers and promoters produce eRNAs and uaRNAs, respectively. Initially proposed as non-functional byproducts of active CREs^16–19^, several studies implicate uaRNAs and eRNAs in cis mechanisms of transcriptional regulation, including RNA polymerase II promoter-proximal pausing^20–22^, recruitment of factors responsible for enhancer-promoter looping^23^, prolonged occupancy of transcription factors (TFs) at CREs^24–26^, and uaRNA- or eRNA-dependent deposition of histone modifications^27,28^. Chromatin-associated RNA mapping has shown that eRNAs associate with target promoters in trans to drive gene activation^25,28,29^. Recently, eRNA-uaRNA duplex formation through complementary ALU elements was shown to contribute to enhancer-promoter looping and, therefore, mRNA regulation^30^. Collectively, this body of literature demonstrates critical functional roles for CRE-associated ncRNAs and, therefore, necessitates novel lines of inquiry into the regulators of uaRNA and eRNA expression.

Many remodelers are recruited to enhancers and promoters to regulate gene expression^7,13,31–43^; however, only a handful have been shown to play a direct role in ncRNA production at CREs^44,45^. esBAF suppresses the production of eRNAs and uaRNAs at target CREs by maintaining the positioning of adjacent nucleosomes^46^. INO80 and BTAF1 directly suppress uaRNA transcription in ES cells^44^, while CHD8 activity at progesterone receptor-dependent enhancers is necessary for eRNA expression^45^. Given the broad diversity of remodelers that localize to CREs in metazoan systems and the potential mechanisms by which they modulate chromatin architecture, we systematically assessed how each of the SNF2-type nucleosome remodeling enzymes contributes to the regulation of the ES cell transcriptome. To that end, we screened all SNF2-type nucleosome remodelers for mRNA, uaRNA, and eRNA regulatory roles in using RNA interference (RNAi) followed by paired steady state and nascent transcriptome profiling to define the contributions of every remodeler to coding and non-coding transcription at CREs. We found that many remodelers regulate thousands of mRNAs, uaRNAs, and eRNAs. Our analyses support previous work showing coordinated regulation between mRNA and eRNA, but not mRNA and uaRNA, transcription. Finally, chromatin-based mechanistic studies suggest two classes of transcription regulation by remodelers: direct regulation through classical mechanisms such as accessibility or transcription factor interactions, and indirect regulation through the maintenance of genomic stability.

## Results

### Screen to define the nucleosome remodeler-regulated ES cell transcriptome

To systematically determine how nucleosome remodelers regulate the transcriptome, we performed an RNAi screen, individually targeting the 32 SNF2-like remodeler ATPases in ES cells (Figure 1A). We depleted the ATPase subunit for each remodeler complex and quantified changes in the coding and non-coding transcriptomes using RNA sequencing (RNA-seq) and Transient Transcriptome sequencing (TT-seq^47^) from the same samples in biological duplicate with high reproducibility (Figure S1). We validated the depletion of 29 remodeler ATPases using RT-qPCR or RNA-seq (Figure 1B, Figure S2A, Table S1). For three ATPases, *Chd6, Smarca1,* and *Ercc6l2*, we could not obtain depletion of 50% or better (as quantified by RT-qPCR) and, therefore, did not proceed further with these candidates.

**Figure 1:**
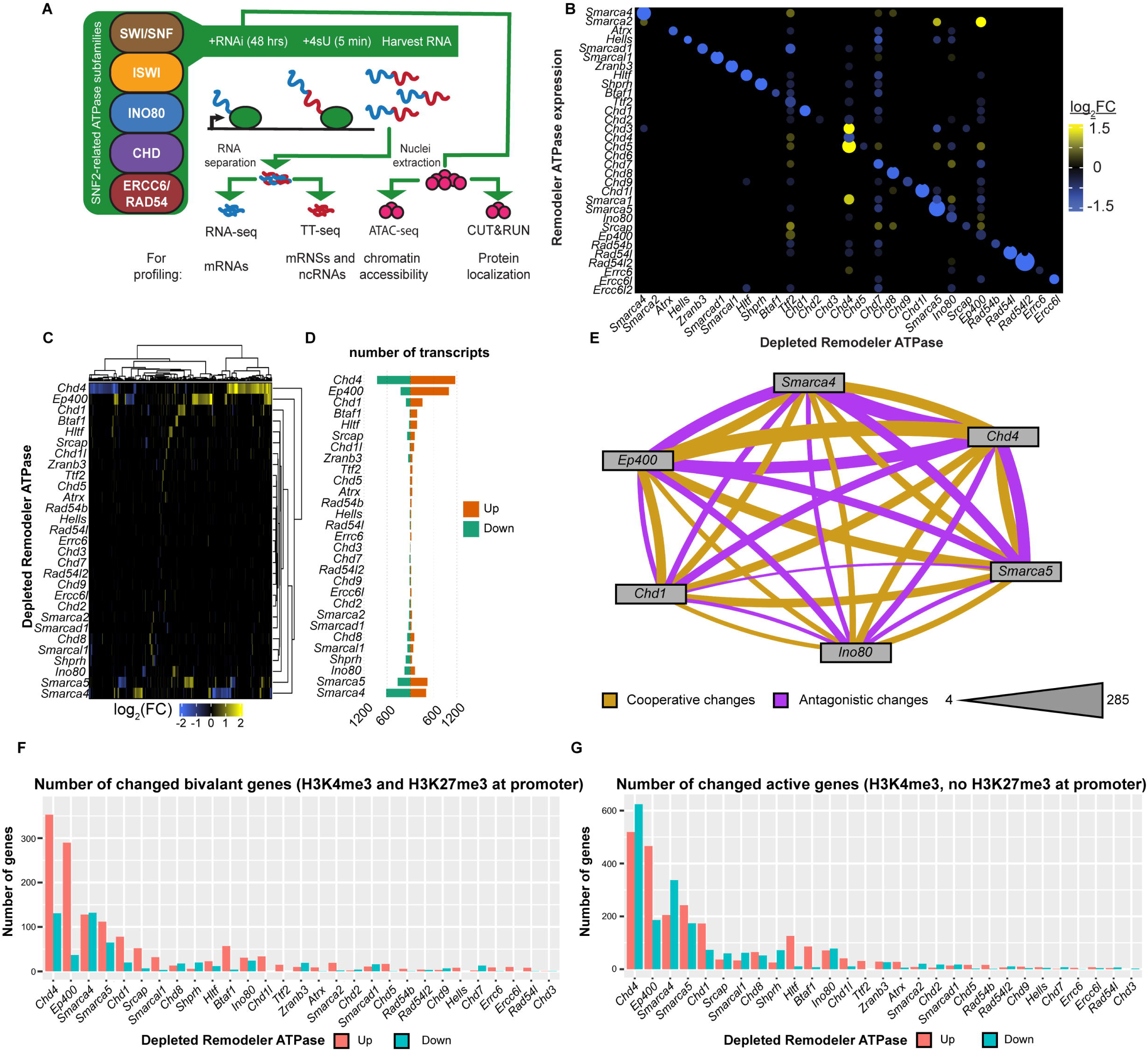
Remodeler ATPases collectively regulate thousands of mRNAs. A. Experimental design for this study. 32 SNF2-related ATPases representing five families were targeted via RNAi in mES cells and the transcriptomic consequences of individual depletions were examined using nascent (TT-seq) and steady state (RNA-seq) approaches. Three ATPases were further examined using genome-wide chromatin accessibility (ATAC-seq) and protein localization (CUT&RUN) profiling. B. Heatmap showing the change in mRNA expression (RNA-seq) of 32 ATPases (y-axis) upon individual depletion of 29 remodelers (x-axis; |log_2_(FC)|≥0.75 and FDR≤0.05). Dot size demonstrates significance. C. Heatmap showing the change in mRNA transcription for genes changed across all TT-seq datasets (|log_2_(FC)|≥0.75 and FDR≤0.05). n=4,406 transcripts. D. Barplot quantifying the number of mRNAs with increased (orange) or decreased (green) transcription upon remodeler depletion using TT-seq (|log_2_(FC)|≥0.75 and FDR≤0.05). E. Network representing the number of mRNAs with altered transcription in the same direction (cooperative, purple) or opposite direction (antagonistic, yellow) shared between six depletion TT-seq datasets (|log_2_(FC)|≥0.75 and FDR≤0.05). Thickness of line represents the number of mRNAs shared between datasets, within the range listed. F. Barplot showing the numbers of mRNAs with increased (orange) or decreased (green) transcription from TT-seq data (|log_2_(FC)|≥0.75 and FDR≤0.05) for promoters displaying bivalent chromatin signatures (H3K4me3 and H3K27me3). G. As in F, for promoters displaying active chromatin signatures (H3K4me3 and no H3K27me3).

### A subset of remodelers regulates the expression of other ATPases in ES cells

To establish remodeler regulation across ATPases, we examined how depletion of a single remodeler ATPase alters the transcript abundance (RNA-seq) of the other 31 SNF2-like ATPases. Most ATPase depletions did not alter the expression of other remodeler ATPases (Figure 1B). However, individual depletion of ten ATPases significantly altered (log_2_(FC)≥0.75 and FDR≤0.05) the mRNA abundance of one or more remodeler ATPases. While depletion of *Ttf2* or *Chd7* altered the expression of many remodeler ATPases (18 and 19, respectively), depletion of *Smarca4*, *Hltf*, *Chd4*, *Chd8*, *Smarca5*, *Ino80*, *Srcap*, or *Ep400* altered the expression of a smaller number of ATPases (between 1 and 14). In some cases, these transcript changes align with known subunit replacements. For example, depletion of *Chd4* led to increases in transcript abundance of *Chd3* and *Chd5*, which serve as alternative ATPases to *Chd4* in the NuRD complex in some cell types^48^ and *Smarca4* (BRG1) depletion leads to increased *Smarca2* (BRM) expression, where BRM is an alternative ATPase for BRG1 in BAF (mSWI/SNF) complexes^49^. Therefore, a subset of remodeler depletions alter the expression of other remodelers and two remodeler depletions, *Ttf2* or *Chd7*, lead to a more global response.

### Nucleosome remodelers collectively regulate the transcription of thousands of mRNAs

To understand how remodelers impact the ES cell mRNA transcriptome, we examined changes in genome-wide mRNA abundance (RNA-seq) and transcription (pre-mRNA levels; TT-seq) upon individual remodeler depletion. We focused on protein-coding features defined in Gencode (detailed in Methods)^50^ and examined 21,596 protein-coding genes, 18,649 of which were actively transcribed (Figure S2B). We defined 5,810 differentially expressed mRNAs across all RNA-seq datasets and 4,406 differentially transcribed mRNAs across all TT-seq datasets (log_2_(FC) ≥0.75 & FDR≤0.05; Figure 1C-D, Figure S2C-D). RNA-seq and TT-seq datasets show similar mRNA profiles apart from five depletions, three of which have been shown to regulate mRNA processing (TTF2^51^) or nascent transcription (CHD1, BTAF1^52–54^; Figure S2E-G).

The largest numbers of mRNAs were differentially expressed upon depletion of *Chd4* (RNA-seq: 2,347, TT-seq: 2,026), *Smarca4* (RNA-seq: 1,099, TT-seq: 1,034), or *Ep400* (RNA-seq: 1,223, TT-seq: 1,251), which encode the ATPases of the NuRD (CHD4), esBAF (BRG1), and Tip60-p400 (p400) complexes, respectively, all of which have been defined as regulators of mRNA expression of ES cells^4,5,55^ (Figure 1C-D, Figure S2C-D). Changes in smaller numbers of transcripts were detected upon depletion of *Smarcad1* (SMARCAD1; RNA-seq: 70, TT-seq: 80)*, Smarca5* (SNF2H; RNA-seq: 553, TT-seq: 772)*, Ino80* (INO80; RNA-seq: 685, TT-seq: 269)*, Srcap* (SRCAP; RNA-seq: 375, TT-seq: 198)*, Chd1* (CHD1; RNA-seq: 16, TT-seq: 428)*, Btaf1* (BTAF1; RNA-seq: 22, TT-seq: 194), or *Chd8* (CHD8; RNA-seq: 154, TT-seq: 185), which encode the ATPases of remodelers previously shown to regulate mRNA transcription in ES cells^44^. We also observed changes in mRNA expression upon depletion of ATPases with less defined roles in transcription, including *Hltf* (HLTF; RNA-seq: 171, TT-seq: 209)*, Zranb3* (ZRANB3; RNA-seq: 10, TT-seq: 112)*, Shprh* (SHPRH; RNA-seq: 110, TT-seq: 164), and *Smarcal1* (SMARCAL1; RNA-seq: 4, TT-seq: 155; Figure 1C-D,Figure S2C-D). Many of these factors have roles in the maintenance of genomic stability^56–59^. Except for *Ttf2* and *Chd7*, depletion of the remaining remodelers displayed approximately 100 or fewer differentially expressed mRNAs (Figure 1C-D, Figure S2C-D).

When examining the specificity of mRNA regulation by remodelers, we found that 55% (2421) of all differentially transcribed mRNAs were changed in only one remodeler depletion indicating that many mRNAs are regulated by only one remodeler (Figure S2H, Table S2). However, 45% of changed mRNAs were changed in 2 or more depletions, reflecting previous reports that remodeler binding at CREs show substantial overlap^42^. To better understand joint regulation of mRNA transcription by remodelers, we examined the pairwise shared (cooperative) or opposite (antagonistic) effects on mRNA transcription between *Ep400*, *Smarca4*, *Chd4*, *Smarca5*, *Chd1*, and *Ino80*, all encoding remodeler ATPases with established roles in pluripotency regulation (Figure 1E)^4,6,7,9,13^. *Smarca4* and *Chd4* displayed the largest number of mRNAs with antagonistic changes, in line with prior studies^60,61^, whereas *Chd4* and *Ep400* had the most overlapping mRNAs with cooperative changes. Toward understanding the regulatory relationships of mRNA transcription between all remodelers, we examined differential expression data across all the TT-seq datasets using principal component analysis (PCA), focusing on mRNAs changed in 3 or more depletions (851 total mRNAs, Figure S2I). PC1-3 accounted for 57% of data variance and were driven by regulatory interactions between *Chd4*, *Ep400*, *Smarca4*, and *Smarca5* (Figure S2I). Overall, these data indicate BRG1, p400, CHD4, and SNF2H are the major drivers of mRNA regulation among remodelers in ES cells, as supported by prior studies, but highlight the contribution of all remodelers to mRNA expression in ES cells.

### Epigenomic analyses suggests roles for BTAF1, CHD8, SRCAP, SMARCAL1, SHPRH, and HLTF in bivalent and active gene regulation in ES cells

To assess whether the local epigenomic environment can categorize mRNA regulation by remodelers, we analyzed available ChIP-seq datasets from WT ES cells^62^ to define mRNA promoters enriched for the post-translational histone modifications H3K4me3 (active genes) or H3K4me3 and H3K27me3 (bivalent genes^63,64^). Depletion of *Smarca4*, *Ep400, Chd4*, *Chd1*, or *Smarca5* led to transcriptional changes in bivalent genes (Figure 1F), consistent with expectations for remodelers with roles in pluripotency^4,5,13,42,65^. These data demonstrate the utility of integrating transcriptomic and epigenetic data to understand remodeler activity, prompting a similar exploration into other remodelers.

We examined the H3K4me3 and H3K27me3 profiles for genes changed in every remodeler depletion. Similar to *Chd4* and *Ep400*, *Srcap* or *Smarcal1* depletion resulted in upregulation of bivalent genes (Figure 1F) suggesting SRCAP and SMARCAL1 contribute to pluripotency through bivalent gene regulation. By contrast, *Chd8* depletion altered transcription of active genes, consistent with previous work showing a poor correlation between CHD8 and H3K27me3 occupancy (Figure 1F-G)^42^. Similarly, depletion of *Shprh* or *Hltf* altered the transcription of genes with active epigenetic profiles (Figure 1F-G). Depletion of *Btaf1* led to the upregulation of bivalent and active genes, consistent with a general role in transcriptional regulation^44^. Therefore, our integrated analyses recapitulate known trends for some remodelers (e.g., CHD4 and p400), support suggested trends for other remodelers (e.g., BTAF1, SRCAP, and CHD8), and suggest as-yet unestablished roles for other remodelers (e.g., SMARCAL1, HLTF, and SHPRH).

### Remodeler depletions collectively alter non-coding transcription at thousands of promoters

Divergent transcription originating from mRNA promoters yields non-coding RNAs termed uaRNAs (also called promoter-associated non-coding RNAs [pancRNAs] or promoter-proximal transcripts [PROMPTs])^66,67^. Previous work has shown that a few remodeler ATPases, including BRG1, INO80, and BTAF1, suppress non-coding transcription from promoters in ES cells^44,46^, with homologs of BAF, CHD1, and SNF2H having similar functions in regulating non-coding transcription within gene bodies in fission and budding yeast^68–70^. We sought to define the remodeler-dependent uaRNA transcriptome using our TT-seq datasets. We detected antisense transcription from 20,615 promoters (Figure S3A). Of these, 2,645 promoters showed significantly altered uaRNA transcription (log_2_(FC)≥0.5 and FDR≤0.05) across all depletions (Figure 2A-B). These altered uaRNAs represent 12.8% of transcribed uaRNAs detected in our datasets and are among the most highly transcribed uaRNAs (Figure S3B). Remodeler depletions altering transcription of the greatest number of mRNAs also altered the transcription of the greatest number of uaRNAs (Figure S3C).

**Figure 2:**
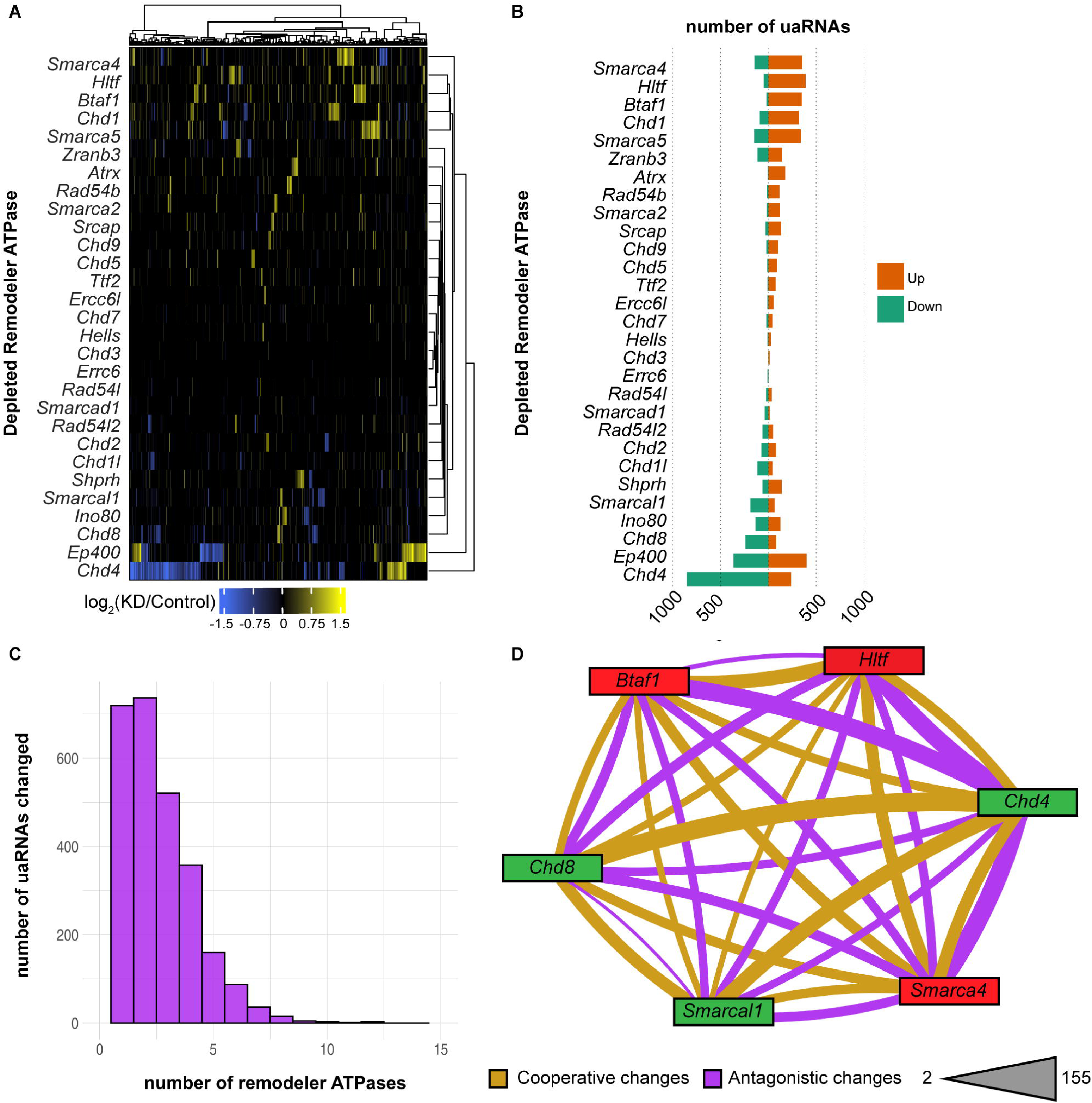
Remodelers contribute to uaRNA transcription regulation in ES cells. A. Heatmap showing the change in transcription of uaRNAs for transcripts changed across all TT-seq datasets (|log_2_(FC)|≥0.5 and FDR≤0.05). n=2,645. B. Barplot quantifying the number of uaRNAs with increased (orange) or decreased (green) transcription in each depletion TT-seq dataset (|log2(FC)|≥0.5 and FDR≤0.05). C. Barplot showing distribution of uaRNAs altered in one or more depletion TT-seq dataset(s) (|log_2_(FC)|≥0.5 and FDR≤0.05). D. Network representing the number of uaRNAs with altered transcription in the same direction (cooperative, purple) or opposite direction (antagonistic, yellow) shared between six depletion datasets (|log_2_(FC)|≥0.5 and FDR≤0.05). Thickness of line represents the number of uaRNAs in category shared between datasets, within the range listed. Green labels indicate “activator” class and red indicate “repressors” class remodelers.

Disruption of remodelers had a range of effects on uaRNA transcription, where depletion of *Smarca4*, *Hltf1, Btaf1*, *Chd1,* or *Smarca5* resulted in mostly upregulation of uaRNAs (repressors), depletion of *Chd4*, *Chd8*, or *Smarcal1* resulted in mostly downregulation of uaRNAs (activators) and depletion of *Ep400*, *Ino80*, *Shprh*, *Chd2*, or *Zranb3* resulted in bidirectional changes to uaRNA transcription (Figure 2A-B, Table S3). These remodeler ATPases or their orthologs localize to promoter elements in metazoan systems^33,42,43,58,71^, suggesting possible direct regulation of uaRNA transcription. Depletion of several other ATPases, including *Atrx*, *Chd1l*, *Chd9*, *Rad54b*, or *Rad54l2*, affected the transcription of between 111 and 258 uaRNAs while the remaining depletions affected less than 100 uaRNAs (Figure 2B, Table S3). When evaluating the specificity of remodelers in uaRNA regulation, we found that 27% of changed uaRNAs were differentially transcribed in only one remodeler depletion (Figure 2C, Table S2), compared to 55% for uniquely regulated mRNAs (Figure S2H, Table S2). These data suggest that uaRNA regulation exhibits more overlapping effects by remodelers compared to remodeler-based mRNA regulation.

Given the emergence of remodelers acting as uaRNA repressors or activators, we examined pairwise cooperative and antagonistic relationships between three repressors (BRG1, HLTF, and BTAF1, red in Figure 2D) and three activators (CHD4, SMARCAL1, and CHD8, green in Figure 2D) on uaRNA transcription. Consistent with these classifications, repressors and activators had more cooperative interactions with other remodelers in the same class, with antagonistic relationships observed between repressors and activators (Figure 2D). We examined the regulatory relationships between all remodelers on uaRNAs differentially transcribed in 3 or more depletions using PCA (n=1,118 uaRNAs). PC1 (24%) was primarily driven by antagonistic relationships between *Chd4* KD and uaRNA repressor remodelers, such as *Smarca4*, *Hltf*, or *Chd1*, and cooperative relationships with other uaRNA activators, such as *Smarcal1* and *Chd8* (Figure S3D). PC2 (14%) was defined by the distinction of *Ep400* from all other remodelers, most likely due to its strong bidirectional effect on uaRNA transcription. PC3 (9%) was driven by the distinctions between *Smarca4*, *Smarca5*, and other uaRNA repressors (Figure S3D). These analyses show that, as in mRNA regulation, BRG1, p400, CHD4, and SNF2H exhibit strong and mostly specific influences on uaRNA transcription in ES cells and identify remodeler classes acting as repressors, activators, or bidirectional regulators of uaRNA transcription.

### Remodeler-based changes in uaRNA transcription are not coordinated with changes in mRNA transcription

Prior studies demonstrate that mRNA and uaRNA transcription levels from the same promoter generally correlate^17,18,72^, suggesting that the regulatory effects of remodelers would impact both transcripts originating from a shared promoter. We hypothesized that changes in uaRNA and mRNA transcription would be correlated upon remodeler depletion, given the transcripts share a common promoter. Surprisingly, we identified little coordinated regulation between mRNA and uaRNA transcription upon every remodeler ATPase depletion. Two such examples are illustrated by *Smarca4* KD and *Smarcal1* KD (Figure 3A,B), where only 49 of 2,003 promoter elements in *Smarca4* KD, and only 1 out of 400 promoter elements in *Smarcal1* KD, exhibited concordant changes in mRNA and uaRNA transcription.

**Figure 3:**
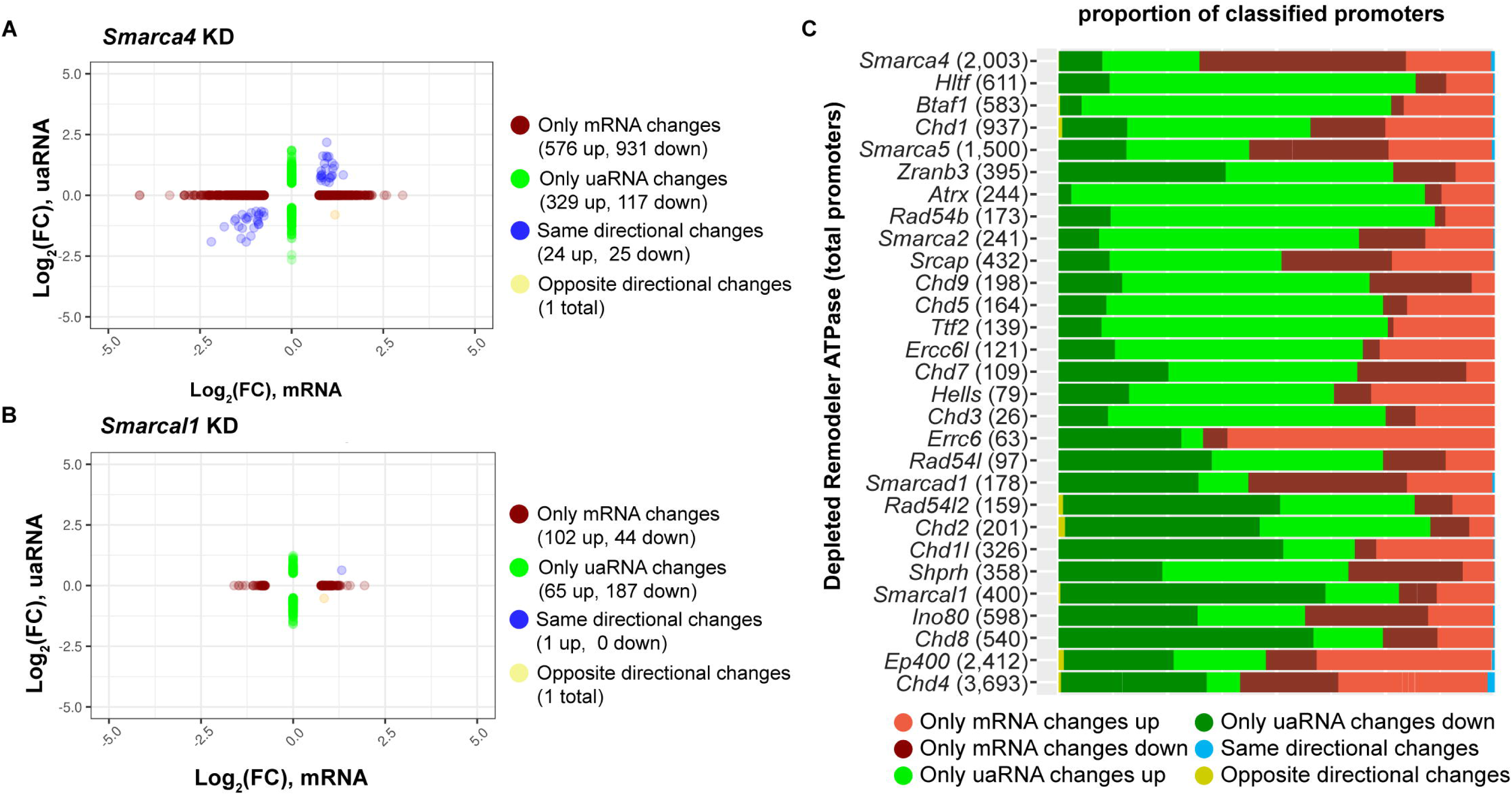
Changes in mRNA and uaRNA transcription are not coordinated in ES cells. A. Scatterplot showing changes in mRNA vs uaRNA transcription in *Smarca4* depletion from TT-seq data. Colors represent promoters sorted into four categories based on mRNA and/or uaRNA change: red = promoters with only significant mRNA changes (mRNA change |log_2_(FC)|≥0.75 and FDR≤0.05, no significant change in uaRNA transcription), green = promoters with only significant uaRNA changes (uaRNA change |log_2_(FC)|≥0.5 and FDR≤0.05, no significant change in mRNA transcription), blue = promoters with significant changes in both transcripts in the same direction (mRNA change |log_2_(FC)|≥0.75 and FDR≤0.05 and uaRNA change |log_2_(FC)|≥0.5 and FDR≤0.05, in the same direction), yellow = promoters with significant changes in both transcripts in opposing direction (mRNA change |log_2_(FC)|≥0.75 and FDR≤0.05 and uaRNA change |log_2_(FC)|≥0.5 and FDR≤0.05, in the opposing direction). B. Same as in A, for *Smarcal1* depletion. C. Barplot displaying the relative distribution of promoters with significant mRNA or uaRNA changed in TT-seq datasets sorted into six categories based on the directional change of each transcript. mRNA change |log_2_(FC)|≥0.75 and FDR≤0.05 and uaRNA change |log_2_(FC)|≥0.5 and FDR≤0.05. Number in parenthesis indicates the total number of promoters showing significant change in mRNA and/or uaRNA change.

To assess the coordination of uaRNA and mRNA transcription regulation by all remodelers, we sorted uaRNA-producing promoter elements into six categories based on mRNA and uaRNA change (mRNA only up, mRNA only down, uaRNA only up, uaRNA only down, same directional change, opposite directional change). We found that for most promoters, remodeler depletion specifically affected only mRNA or uaRNA transcription (Figure 3C) with concordant or discordant directional changes comprising less than 5% of all affected promoters (e.g., *Chd2* or *Chd4* depletion; Figure 3C). Consistent with this, the same directional change and opposite directional change categories were significantly depleted in all remodeler depletions (*P*≤0.05, Chi-squared test of homogeneity assuming equal distribution across all categories, Figure S3E). When comparing promoter directionality in these categories for each remodeler depletion relative to control, we found shifts in directionality consistent with changes in one transcript but not the other (exemplified in *Smarca4* KD and *Smarca5* KD, Figure S3F,G). These data demonstrate that the impact of remodeler depletion on mRNA or uaRNA regulation is not coordinated, further suggesting that mRNA and uaRNA transcription are not co-regulated by remodelers.

Two potential explanations for these findings are that promoters showing only mRNA change do not express uaRNAs or these uaRNAs are not detectable by TT-seq. To evaluate these possibilities, we quantified uaRNA transcription from promoters showing only mRNA or uaRNA changes for a subset of 12 remodelers with at least 50 changed promoters in each group. We found that non-coding transcription could be measured at promoters showing only mRNA change (Figure S3H,I). Therefore, while it remains unclear if our findings extend to promoters with lowly transcribed/undetectable uaRNAs, these data suggest the effects of remodeler depletion on mRNA and uaRNA transcription are largely independent.

### Remodelers broadly regulate eRNA transcription

Another major class of CRE-associated ncRNAs are enhancer RNAs (eRNAs). To understand how remodelers influence non-coding transcription from enhancer elements in ES cells, we annotated 101,587 putative enhancer elements from previously defined TSS-distal DNase I hypersensitive sites (DHSs^73^). From this dataset, we detected transcription from 75,490 TSS-distal DHSs using TT-seq (Figure S4A), which, moving forward, we define as putative enhancers/eRNAs. Across all remodeler depletions, we detected 9,932 putative eRNAs (representing 11.6% of all detected putative eRNAs) with changed transcription (log_2_(FC)≥0.5 and FDR≤0.05) in at least one remodeler depletion (Figure 4A,B, Figure S4A). These changed putative eRNAs are a subset of the most highly transcribed putative eRNAs detected (Figure S4B).

**Figure 4:**
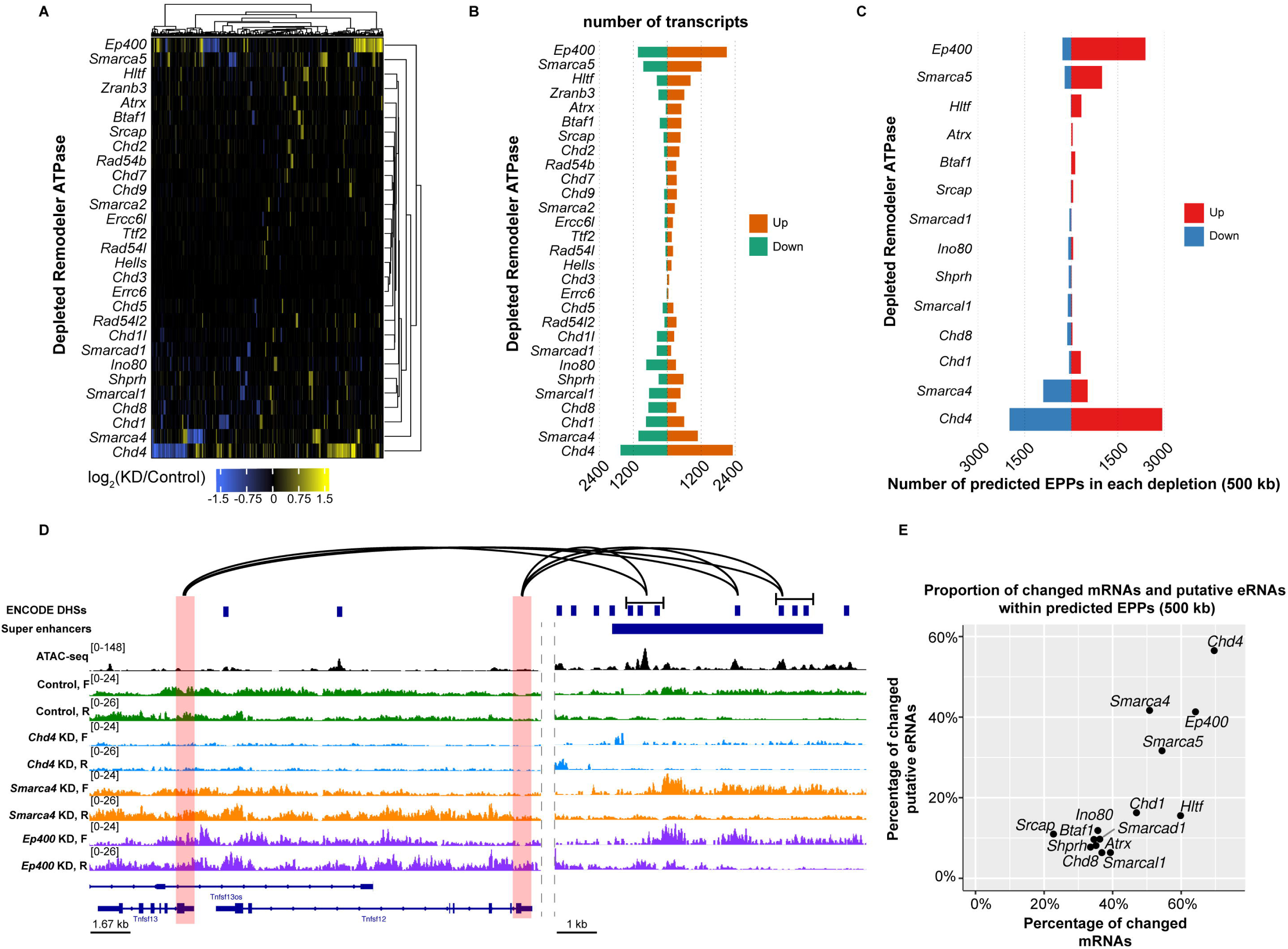
Coordinated regulation of mRNA and enhancer transcription in remodeler depletions. A. Heatmap showing the change in transcription of putative enhancers (|log_2_(FC)|≥0.5 and FDR≤0.05) across TT-seq datasets. n=9,932 regions. B. Barplot showing the total number of putative enhancers with increased (orange) or decreased (green) transcription in each depletion TT-seq dataset (|log2(FC)|≥0.5 and FDR≤0.05). C. Barplot showing the number of predicted enhancer-promoter pairs (EPPs) with increased or decreased transcription in each TT-seq dataset based on a 500 kb distance threshold. D. Browser track showing chromatin accessibility (ATAC-seq) and comparing transcription (TT-seq) in the control, *Chd4* KD, *Smarca4* KD, and *Ep400* KD over the *Tnfsf12* locus and the interacting super-enhancer region. Red boxes highlight promoter regions of each locus, and lines indicate DHSs within the super enhancer region shown to interact with each promoter as shown using available promoter capture HiC data^78–80^. E. Scatterplot showing the relative proportions of significantly changed mRNAs and putative eRNAs that could be placed into predicted EPPs in each depletion dataset based on a 500 kb distance threshold.

Consistent with our findings on remodeler regulation of mRNA and uaRNA transcription, depletion of *Chd4*, *Ep400*, *Smarca5*, or *Smarca4* led to the largest numbers of putative eRNAs with changed transcription (4,314, 3,412, 2,240, and 2,277 changed putative eRNAs, respectively; Figure 4A,B, Table S3). Depletion of other remodelers important for eRNA transcription in ES cells or other systems, including *Ino80*, *Chd8*, and *Chd1*^36,45,74^, showed robust changes in putative eRNA transcription as well (1,139, 1,070, and 1,469 changed putative eRNAs, respectively; Figure 4A,B, Table S3). Beyond these previously characterized regulators, we observed changes in putative eRNA transcription in every remodeler depletion, with two remodeler depletions changing the transcription of thousands of putative eRNAs (*Hltf* KD: 1,292 and *Smarcal1* KD: 1,210) and the remaining altering transcription of between 71 and 995 putative eRNAs (Figure 4A,B, Table S3). Further, the number of putative eRNAs changed was proportional to the number of mRNAs changed upon each remodeler depletion (Spearman’s rho=0.925; Figure S4C). When evaluating the specificity of putative eRNAs, we found that 78% of changed putative eRNAs (7,479) were differentially transcribed in two or more remodeler depletions (Figure S5D). Therefore, our data reveal that perturbation of many SNF2-related ATPases alter eRNA transcription in ES cells.

We next examined the pairwise regulatory relationships in putative eRNA regulation with the six remodeler ATPase depletions that had the largest effects: *Chd4*, *Ep400*, *Smarca4*, *Smarca5*, *Hltf* and *Smarcal1.* We found that *Smarcal1* displayed relatively balanced cooperative/antagonistic relationships with *Smarca4*, *Chd4*, and *Smarca5*, but a stronger antagonistic relationship with *Ep400* (Figure S4E). On the other hand, *Hltf* showed balanced cooperative/antagonistic relationships with *Smarca4*, *Chd4*, and *Ep400*, and a strong cooperative relationship with *Smarca5*. These data show specificity in the regulatory interactions with these two understudied remodelers and previously established regulators of transcription in ES cells. We further probed the regulatory interactions among all remodelers through PCA using differential expression data focused on putative eRNAs changed in 3 or more depletions (n=4,882 putative eRNAs; Figure S4F). PC1-3 account for 48% of all the variance and, similar to the mRNA data, were defined by the regulatory interactions of *Chd4*, *Ep400*, *Smarca4*, and *Smarca5* (Figure S4F). Together, these data demonstrate broad regulation of putative eRNA transcription by remodelers, with the same four remodelers that have the greatest contribution to the mRNA transcriptome having the largest impact on the eRNA transcriptome. Furthermore, we have identified SMARCAL1 and HLTF as novel regulators of eRNA transcription.

### Remodeler depletions lead to coordinated changes in putative eRNA and mRNA transcription

Current models suggest transcriptional changes from functionally associated enhancers and mRNAs correlate with one another^75^, leading us to hypothesize that putative enhancer-promoter pairs (EPPs) would show correlated changes within our depletion TT-seq datasets. We performed an *in silico* search for pairs of enhancers and mRNA promoters showing correlated transcriptional change within our TT-seq data across multiple distance thresholds (100 kb, 250 kb, 500 kb, and 1 Mb; see Methods). Consistent with our hypothesis, we detected significantly more putative EPPs than expected by chance across 3 or more distance thresholds for 14 remodelers (Benjamini & Yekutieli padj values≤0.05, Figure S5A,B). At the 500 kb distance threshold, the median distance between EPPs within each depletion at this threshold ranged from 96-225 kb (Figure S5C), well within the range of functionally validated EPPs in mammalian systems^76,77^. Included in remodeler depletions that displayed significant enrichment for EPPs were known or suspected regulators of enhancer activity, including CHD4, BRG1, SNF2H, p400, CHD1, INO80, and CHD8. Several other remodelers behaved similarly to (or had a greater enrichment than) these known enhancer regulators, including HLTF, SHPRH, SMARCAL1, BTAF1, SRCAP, SMARCAD1, and ATRX (Figure S5A,B). These analyses suggest that depletion of at least 14 remodelers induce coordinated mRNA and putative eRNA transcriptional changes. We selected the 500 kb distance threshold for further characterization of EPPs.

*Chd4*, *Ep400*, *Smarca5*, and *Smarca4* depletions displayed the largest numbers of changed EPPs (Figure 4C). Analyzing published promoter capture HiC data^78–80^, predicted EPPs include correlated transcriptional changes between super-enhancers and nearby target genes, supporting these analyses identified functional interactions (Figure 4C, Figure S5D). We found that 23-69% of changed mRNAs and 10-60% of putative eRNAs could be assigned to EPPs across these 14 depletions (Figure 4E). To understand the biological pathways related to the EPPs in our datasets, we performed gene ontology (GO) analyses on changed mRNAs in upregulated and downregulated EPPs in each of the 14 depletions. Consistent with roles in pluripotency regulation, mRNAs in upregulated EPPs in *Chd4*, *Ep400*, and *Srcap* KD were enriched for terms related to differentiation (Figure S5E). DNA replication-dependent histone genes were abundant within EPPs upregulated upon *Btaf1* or *Hltf* KD and downregulated by *Chd4*, *Chd8*, or *Smarcal1* KD, contributing to enrichment of chromatin assembly or nucleosome organization GO terms (Figure S5E,F). These analyses suggest coordinated changes in putative eRNA and mRNA transcription upon depletion of at least 14 remodelers, with specifically enriched biological processes.

### CHD8 and SRCAP, but not SMARCAL1, co-localize with transcriptomic changes in ES cells

To mechanistically determine how a subset of these remodelers are acting in ES cells to regulate the transcriptome, we selected three ATPases for further characterization: CHD8, SRCAP, and SMARCAL1. First, we examined the chromatin localization of these three remodelers using Cleavage Under Target and Release Using Nuclease (CUT&RUN) for SRCAP and SMARCAL1 and analyzing available ChIP-seq data from ES cells for CHD8^42^. We defined 50,743, 30,053, and 2,271 peaks for SRCAP, CHD8, and SMARCAL1, respectively (Figure 5A,B, Figure S6A). To examine whether these factors may be acting directly at locations of transcription changes, we integrated our transcriptomic and localization data for each remodeler. We found that CHD8 and SRCAP bound many of the promoters and enhancers with transcription changes upon depletion of these remodelers (Figure 5C,D). We observed minimal SMARCAL1 enrichment at changed CREs, suggesting indirect effects of SMARCAL1 loss on transcription (Figure S6B). These data suggest that CHD8 and SRCAP are acting directly to regulate transcription while SMARCAL1 may act through indirect mechanisms.

**Figure 5:**
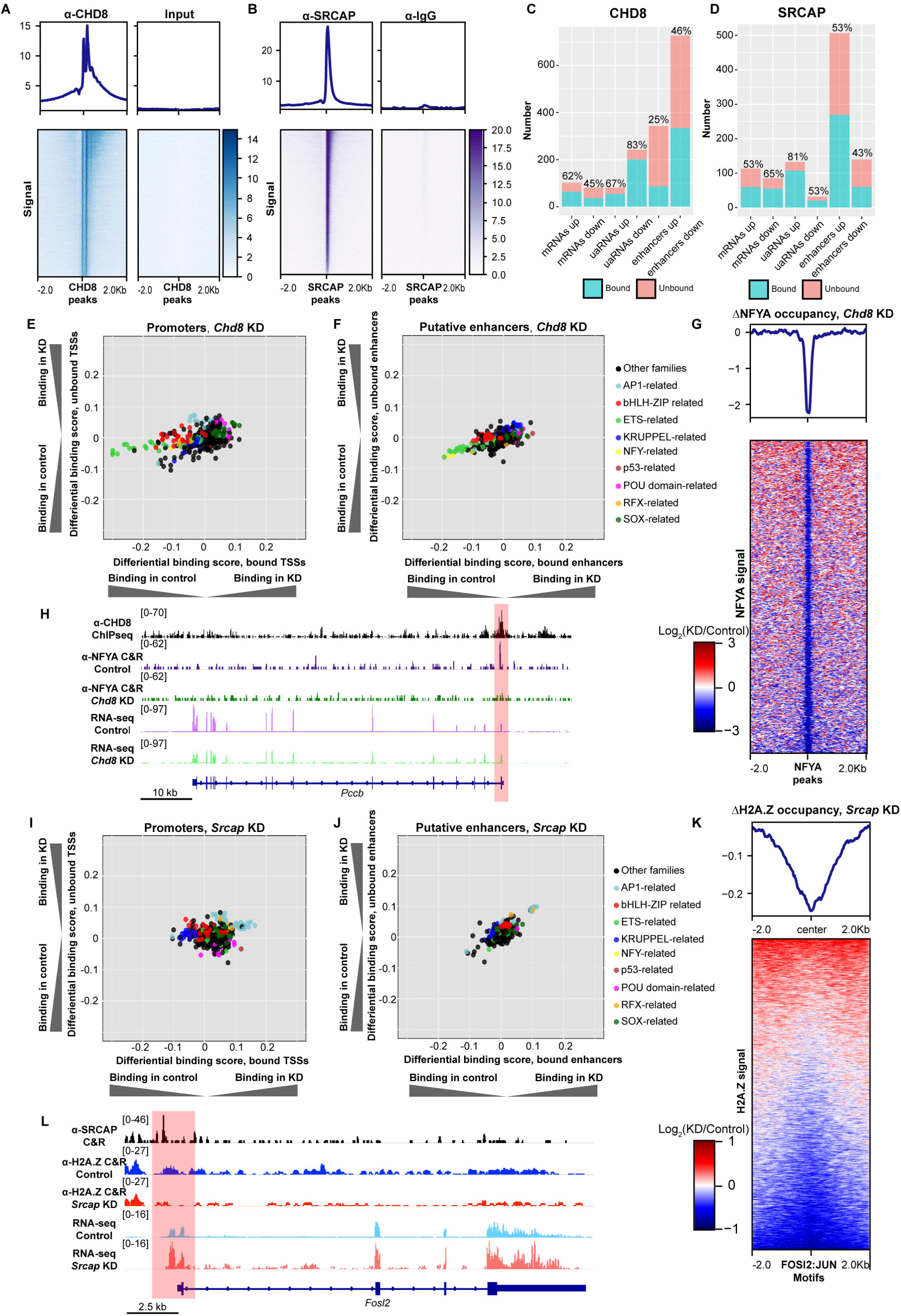
CHD8 and SRCAP regulate transcription through TF-based mechanisms. A. Heatmap showing the enrichment of CHD8 binding over called peaks relative to input from available ChIP-seq data (GSE64825^42^). n=30,053 peaks. B. Heatmap showing SRCAP binding over called peaks relative to IgG using CUT&RUN. n=50,743 peaks. C. Barplot showing the number of CREs within each altered transcript class bound by CHD8 using available ChIP-seq data (GSE64825^42^). Blue represents number of bound elements and pink indicates number of unbound elements. D. As in C, for SRCAP binding using CUT&RUN. E. Scatterplot comparing the binding score of different TF motifs at promoters bound versus not bound by CHD8. Motifs representing factors from similar related groups are colored according to the legend. All factors shown have *P*≤0.05 at bound and unbound loci, defined from ATAC-seq data using TOBIAS. F. As in D, for putative enhancers bound and unbound by CHD8. G. Heatmap showing the change in NFYA binding at NFYA peaks in the *Chd8* KD relative to control CUT&RUN experiments. n=1,179 peaks. H. Browser track showing chromatin accessibility (ATAC-seq) and comparing NFYA binding (CUT&RUN) and gene expression (RNA-seq) in the control and *Chd8* KD at the *Pccb* locus. Red box indicates the promoter region. I. As in E, for promoters bound and unbound by SRCAP. J. As in F, for putative enhancers bound and unbound by SRCAP. K. Heatmap showing the change in H2A.Z binding over FOSL2:JUN motifs defined by JASPAR in the *Srcap* KD relative to control CUT&RUN experiments. n=28,392 loci. L. Browser track showing chromatin accessibility (ATAC-seq) and comparing H2A.Z localization (CUT&RUN) and gene expression (RNA-seq) in the control and *Srcap* KD at the *Fosl2* locus, which was differentially transcribed in the *Srcap* depletion (log_2_(FC)=1.04 and FDR=3.3×10^-7^. Red box indicates the promoter region.

### Transient depletion of *Chd8*, *Srcap*, or *Smarcal1* result in moderate changes to chromatin accessibility and H3K27ac enrichment

To determine whether a chromatin-based mechanism could explain the transcriptomic changes observed upon depletion of CHD8, SRCAP, or SMARCAL1, we examined changes in chromatin accessibility using Assay for Transposon Accessible Chromatin sequencing (ATAC-seq) and H3K27ac enrichment using CUT&RUN following depletion of *Chd8*, *Srcap*, or *Smarcal1.* We identified only 376, 686, and 315 regions with differential accessibility upon depletion of *Chd8*, *Srcap*, and *Smarcal1*, respectively, with minimal overlap of CREs with changes in transcription (Figure S6C). We detected 21, 321, and 211 differentially enriched H3K27ac regions upon depletion of *Chd8*, *Srcap,* and *Smarcal1*, respectively, with little overlap with changes in transcription (Figure S6D). These data suggest alternative mechanisms of transcription regulation by CHD8, SRCAP, and SMARCAL1.

### CHD8 regulates binding of NFY-, ETS-, and p53 family TFs

To test whether depletion of *Chd8*, *Srcap*, or *Smarcal1* may influence TF binding, we integrated ATPase localization and ATAC-seq data upon remodeler depletion to analyze TF footprints at promoters and putative enhancers bound by these factors, using Transcription factor Occupancy prediction By Investigation of ATAC-seq Signal (TOBIAS)^81^. TOBIAS compares accessibility over TF motifs to infer differences in TF binding between experimental conditions, providing a proxy of how TF binding profiles change between conditions. Upon *Chd8* depletion, we observed a strong and specific decrease in predicted binding of ETS- and NFY-related factors and a moderate increase in predicted binding of p53-related factors at both promoters and putative enhancers, with the strongest effect at CREs bound by CHD8 (Figure 5E,F). CHD8 has been reported as a positive regulator of ETS-family factor binding in human neurons^82^ and a negative regulator of p53 binding in ES cells^8,83^, both validating our approach and suggesting a conserved role between CHD8 and ETS-factors in ES cells. The association between NFYA binding and CHD8 represents a novel regulatory interaction. Therefore, we assessed changes in NFYA localization in *Chd8* depleted ES cells relative to control using CUT&RUN. We observed a decrease in NFYA binding in *Chd8* depleted ES cells, validating our findings using TOBIAS (Figure 5G). Changes in NFYA binding at promoters bound by CHD8 were associated with transcriptional changes, as exemplified with *Pccb* (Figure 5H). In *Chd8* depleted ES cells, we defined 33 gained and 96 lost NFYA peaks relative to control ES cells (log_2_(FC)≥0.5 and FDR≤0.05, Figure S6E). Using an available ChromHMM map of the ES cell genome^84^, we found that lost peaks were significantly enriched for “Active Promoter” and “Strong Enhancer” chromatin states, while gained peaks were significantly enriched for “Active promoter”,“Intergenic”, and “Weak enhancer” (Fisher’s test, Bonferroni adjusted P value ≤0.05; Figure S6F-G). Notably, NFYA-, ETS-, or p53-family factors were not differentially expressed in *Chd8* depleted ES cells (Table S1), supporting a regulatory interaction between CHD8 and these TFs rather than upstream transcriptional regulation by CHD8. In summary, we have discovered that proper NFYA localization is dependent upon CHD8 in ES cells, and our analyses suggest that CHD8 regulates transcription through functional interactions with ETS, p53, and NFY-family TFs.

### SRCAP drives expression of AP1-related factors through appropriate H2AZ localization

SRCAP and Tip60-p400 are the two remodelers known to incorporate histone variant H2A.Z into nucleosomes ^43,85,86^. Upon depletion of *Srcap*, we found a specific increase in predicted binding of AP1-and RFX-related family factors at promoters and putative enhancers (Figure 5I,J). Loss of H2A.Z localization is associated with increased binding of c-FOS (an AP1 family factor) at promoters in human epithelial cells^87^ and increased FOS expression in mouse embryonic fibroblasts^88^, suggesting that SRCAP also negatively regulates AP1 TF binding in ES cells, perhaps through appropriate H2A.Z incorporation. To test this hypothesis, we determined H2A.Z localization in *Srcap* depleted and control ES cells, finding that *Srcap* depletion results in decreased H2A.Z localization (Figure S6H), as expected given previously defined roles for SRCAP^43^. In line with our hypothesis, we observed depletion of H2AZ localization over FOSL2:JUN motifs in *Srcap* depleted cells (Figure 5K). In support of direct transcriptional regulation of AP1 TFs by SRCAP, AP1 family TFs *Jun* (log_2_(FC)=0.71 and FDR=0.04) and *Fosl2* (log_2_(FC)=1.04 and FDR=3.3×10^-7^) were differentially expressed via RNA-seq in *Srcap* depleted ES cells. SRCAP binds the *Fosl2* promoter in wild-type cells, and, upon *Srcap* depletion, H2A.Z is lost, possibly leading to the increase in transcription observed (Figure 5L). Differential peak analysis identified 121 gained and 735 lost H2A.Z peaks in *Srcap* depleted cells with distribution across enhancer and promoter chromatin states (Figure S6I-K). Together, these data support a mechanism whereby SRCAP contributes to transcriptional regulation of putative eRNAs and mRNAs through maintenance of H2A.Z genomic distribution and direct repression of AP1 TF expression.

In contrast to CHD8 and SRCAP, TOBIAS analysis predicts several TF families with altered binding upon depletion of SMARCAL1 at promoters and enhancers (Figure S6L,M). However, given the small proportion of CREs bound by SMARCAL1 CREs and none of the TOBIAS-identified factors are differentially expressed in *Smarcal1* depleted cells, we did not pursue any candidate TFs.

### Regulatory interactions with Integrator complex may explain changes in ncRNA transcription for remodelers related to genomic stability

In metazoans, post-transcriptional regulation of non-polyadenylated RNAs (including many eRNAs and uaRNAs) occurs at the level of transcription termination through the Integrator complex, which downregulates uaRNA and eRNA biogenesis through cleavage of the nascent RNA and de-stabilization of RNA polymerase II^89–92^. In ES cells, depletion of INTS11 (a component of the cleavage module of Integrator) is associated with a global change in eRNA and uaRNA transcription^93^. Based on these data, we hypothesized that altered binding of Integrator may explain changes in ncRNA transcription upon depletion for remodelers where a direct chromatin-based or TF-based mechanism was not obvious, such as for SMARCAL1. To test this hypothesis, we determined the localization of INTS5, a structural subunit of the Integrator complex, in *Smarcal1* depleted and control ES cells using CUT&RUN. We detected an increase in INTS5 occupancy at promoters and putative enhancers in *Smarcal1* depleted cells (Figure 6A,B).

**Figure 6:**
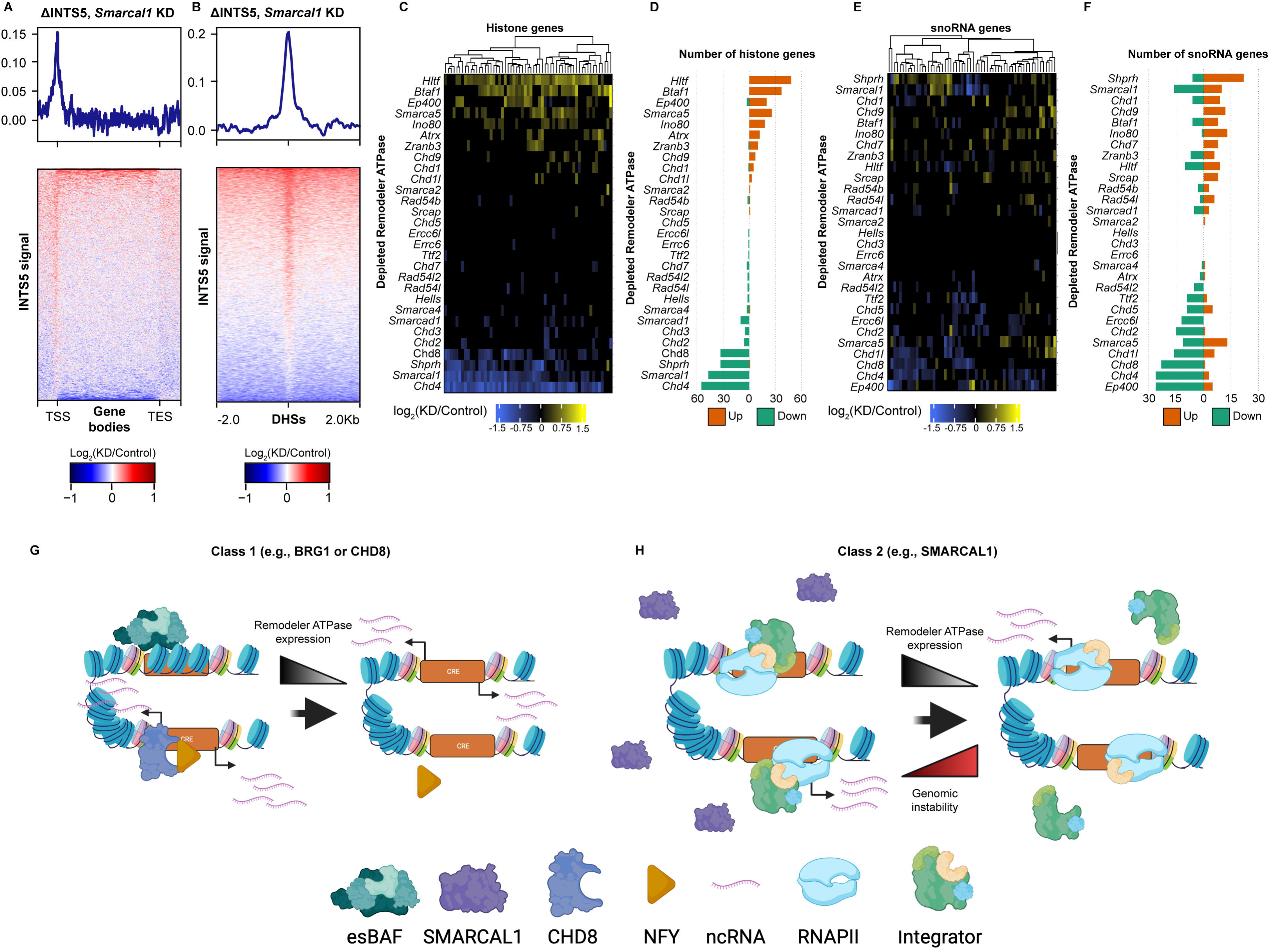
Remodeler depletion results in regulatory interactions with Integrator. A. Heatmap showing the change in INTS5 localization (CUT&RUN) over mRNA gene bodies in *Smarcal1* KD relative to control. n=22,598 loci. B. As in A, over putative enhancers. n=101,588 loci. C. Heatmap showing the change in transcription of histone genes across all 29 TT-seq datasets (FDR≤0.05). n=59. D. Barplot quantifying the number of uaRNAs with increased (orange) or decreased (green) transcription in each depletion TT-seq dataset (FDR≤0.05). E. As in C for snoRNA genes. n=60. F. As in D for snoRNA genes. G. Model depicting class 1 mechanisms of ncRNA regulation by remodelers. H. Model depicting class 2 mechanisms of ncRNA regulation by remodelers.

Given these findings, we examined whether impaired Integrator function could be detected in remodeler depletions. In metazoans, Integrator is critical for proper transcription and 3’ end processing of replication-dependent histone genes and snoRNAs^89,90,94^. To infer Integrator activity, we examined changes in nascent transcription (from TT-seq) of these gene sets in every remodeler depletion. Individual depletion of ten remodelers led to strong upregulation of histone mRNA transcription while depletion of seven remodelers led to a downregulation of these genes (FDR≤0.05, Figure 6C,D), reflecting enriched GO terms such as chromatin assembly or nucleosome organization for mRNAs in predicted EPPs in these depletions as well (Figure S5E,F). Notably, several remodelers that were identified as uaRNA repressors in our screen (e.g., HLTF, BTAF1, SNF2H; Figure 2) led to upregulation of histone transcription, while remodelers identified as uaRNA activators (e.g., CHD4, SMARCAL1, SHPRH; Figure 2) led to downregulation of histone transcription. These transcriptional changes were also observed for snoRNA transcription (FDR≤0.05,Figure 6E,F). In summary, these results support a model wherein changes in Integrator recruitment, assembly, and/or function may contribute to the changes in ncRNA transcription detected at promoters and enhancers upon depletion of some nucleosome remodelers. We propose a model where remodelers act in two classes: Class 1 regulate coding and non-coding transcription through chromatin and/or TF-based mechanisms and Class 2, often remodelers with roles in genome stability, safeguard the genome through diverse activities that reinforce appropriate Integrator localization and ncRNA transcription (Figure 6G,H).

## Discussion

While a handful of the 32 SNF2-related ATPases have been demonstrated as direct transcriptional regulators, many more have been proposed to fulfill this role. Leveraging the well-studied murine ES cell model, we systematically examined the transcriptional consequences to CREs upon individual depletion of 29 SNF2-related ATPases. Our findings support the regulatory paradigm that CHD4, BRG1, p400, and SNF2H act as key contributors to the mRNA transcriptome in ES cells, as demonstrated in previous studies^4,5,9,41,42,95^, with more modest, but still critical, contributions from CHD1, INO80, CHD8, SRCAP, and BTAF1^7,8,13,43,44,53^ (Figure 1). Our study also demonstrates that multiple additional remodelers have modest contributions to mRNA transcriptional regulation in ES cells, including SMARCAL1, HLTF, and SHPRH, consistent with prior work where loss of each of these ATPases has been associated with effects on mRNA transcription in other systems^33,35,40,59,96,97^.

Our analyses suggest that this regulatory paradigm applies to both the coding and CRE-associated non-coding transcriptome, with some notable distinctions. We observed altered transcription in all three classes of transcripts analyzed (mRNAs, uaRNAs, and eRNAs) in each depletion dataset, with the overall number of transcripts changed in each depletion remaining relatively proportional across all three classes. However, while 55% of mRNAs were significantly changed in only one remodeler depletion, only 27% of uaRNAs and 22% of putative eRNAs were changed in only one depletion. These findings suggest that remodelers are more collaborative in the regulation of the non-coding transcriptome.

Our data support BRG1 and BTAF1 acting as repressors of uaRNA transcription, and suggest that several other factors, including HLTF, CHD1, and SNF2H, share in the repression of uaRNA in ES cells (Figure 2). We also observed remodelers operate as activators of uaRNA transcription, most notably CHD4, CHD8, and SMARCAL1. The predominant hypothesis for uaRNA function is currently that they contribute to the regulation of the corresponding mRNA. However, our findings (Figure 3) together with previous studies^44,46,72,98,99^, indicate regulation of the shared mRNA seems unlikely. Rather, we observe that uaRNAs are uniquely and independently regulated from the mRNAs for which they share a promoter. Given the lack of overlap in shared uaRNAs and mRNAs affected upon remodeler depletion, these data also suggest that uaRNAs do not contribute to mRNA transcription.

Our analyses identified changes in enhancer transcription for many ATPase depletions, and we found that at least 14 depletions have corresponding changes in mRNA transcription (Figure 4). We have two non-mutually exclusive explanations for why our analyses predicted EPPs in only 14 remodeler depletions. First, many genes are likely regulated by multiple enhancers and we were only able to detect transcriptional changes in mRNAs where the sole enhancer, primary enhancer, and/or multiple enhancers regulating the gene in question were perturbed by ATPase depletion. Second, the elicited change in eRNA transcription from remodeler depletion (or the corresponding change in mRNA transcription due to elicited change in enhancer activity) was below our stringent significance threshold. We were able to predict statistically significant numbers of EPPs across most distance thresholds in these 14 depletions, with some of these predictions including established super-enhancer and promoter interactions, suggesting this trend is robust. While further studies are needed to validate our findings, our data suggest that depletion of several ATPases induces mRNA transcription associated with changes in enhancer activity in ES cells, including novel contributors SRCAP, HLTF, SMARCAL1, BTAF1, SHPRH, and SMARCAD1.

Our study demonstrates direct mechanisms of action for CHD8 and SRCAP, but an indirect mechanism for SMARCAL1, in CRE-associated transcription regulation (Figures 5 and 6). Depletion of all three remodelers induced moderate changes in chromatin accessibility and H3K27ac distribution where these modest changes were not correlated with transcriptional changes at enhancers or promoters. Our data indicate that CHD8 orchestrates transcriptional changes in ES cells through regulation of TFs, including ETS, p53, and NFY factors (Figure 5). Previously, homozygous deletion of CHD8 was associated with changes in chromatin accessibility in ES cells whereas heterozygous deletion induced almost no detectable changes^8^. As a fraction of functional protein remains in the cell after our depletion approach, it is possible that we have not captured the full effects of this ATPase on chromatin accessibility. Another possible contributing factor is compensation by major regulators of accessibility such as CHD4, BRG1, p400, and SNF2H, as has been recently demonstrated between mSWI/SNF and Tip60/p400 complexes^95^.

Our data support SRCAP contributing to transcriptome regulation through H2A.Z and AP1 factors (Figure 5). It was interesting that over 50% of H2A.Z peaks lost in ES cells upon depletion of *Srcap* were classified as “Strong Enhancer” or “Enhancer”, as the other major regulator of H2A.Z incorporation, Tip60-p400, primarily localizes to promoters in ES cells^4,42^. This finding suggests a functional distinction between these two complexes, where Tip60-p400 targets H2A.Z to promoters while SRCAP loads H2A.Z at enhancers in ES cells. Our data also show that loss of SRCAP results in a redistribution of H2A.Z to bivalent and repressed chromatin, indicating that SRCAP contributes to pluripotency in ES cells through maintenance of proper H2A.Z localization. For both CHD8 and SRCAP, future studies using rapid depletion approaches such as the dTAG or PROTAC systems would help to inform their direct influences on the transcriptome^100,101^.

While our studies did not support direct action of SMARCAL1 in CRE-associated transcription regulation, the observed increase in INTS5 localization at promoters and enhancers upon *Smarcal1* depletion suggests a change in post-transcriptional regulation accounts for the transcriptional changes observed (Figure 6). The Integrator complex is a multifaceted complex with roles in transcription termination, replication dependent histone gene expression, snoRNA processing, and genomic stability^89,90^. Stressful growth conditions trigger genome-wide transcription termination defects^102,103^ (similar to what has been seen upon rapid depletion of INTS11 in mouse ES cells^93^), de-regulation of uaRNA transcription^67^, and a decrease in interactions between Integrator complex subunits and RNA polymerase II^104^. Given these previous findings and our findings regarding changes in Integrator subunit localization and widespread changes in uaRNA, eRNA, histone, and snoRNA transcription, we propose that the transcriptomic effects observed under depletion of ATPases involved in genomic stability, such as SMARCAL1 and others, may be attributed, in part, to defective Integrator recruitment, binding, and/or activity. Transcriptional and epigenetic defects as a consequence of genomic instability are a well-documented phenomenon^105–109^, and when considering that many SNF2-like ATPases are critical DNA repair factors, this provides a plausible mechanism to explain some of the transcriptomic effects we observe. Therefore, we propose that remodelers can influence non-coding transcription at CREs through two major mechanistic classes. Class 1 includes the traditional mechanisms of regulation, such as regulation of chromatin accessibility, nucleosome positioning (such as BRG1) or TF occupancy (such as CHD8; Figure 6G). Class 2 includes indirect mechanisms of action, centered around maintenance of genomic stability to limit stress responses and ensure transcriptional fidelity (such as SMARCAL1; Figure 6H). Importantly, as many remodelers have described roles in transcription and genomic stability, these classes are non-mutually exclusive. Further, we observe examples of upregulation and downregulation of non-polyadenylated RNAs in several remodeler depletions, suggesting that these ATPases may influence Integrator assembly or activity in different ways. Future studies will be needed to understand the mechanism through which these factors influence Integrator localization, assembly, and activity.

---

**Figure S1:**
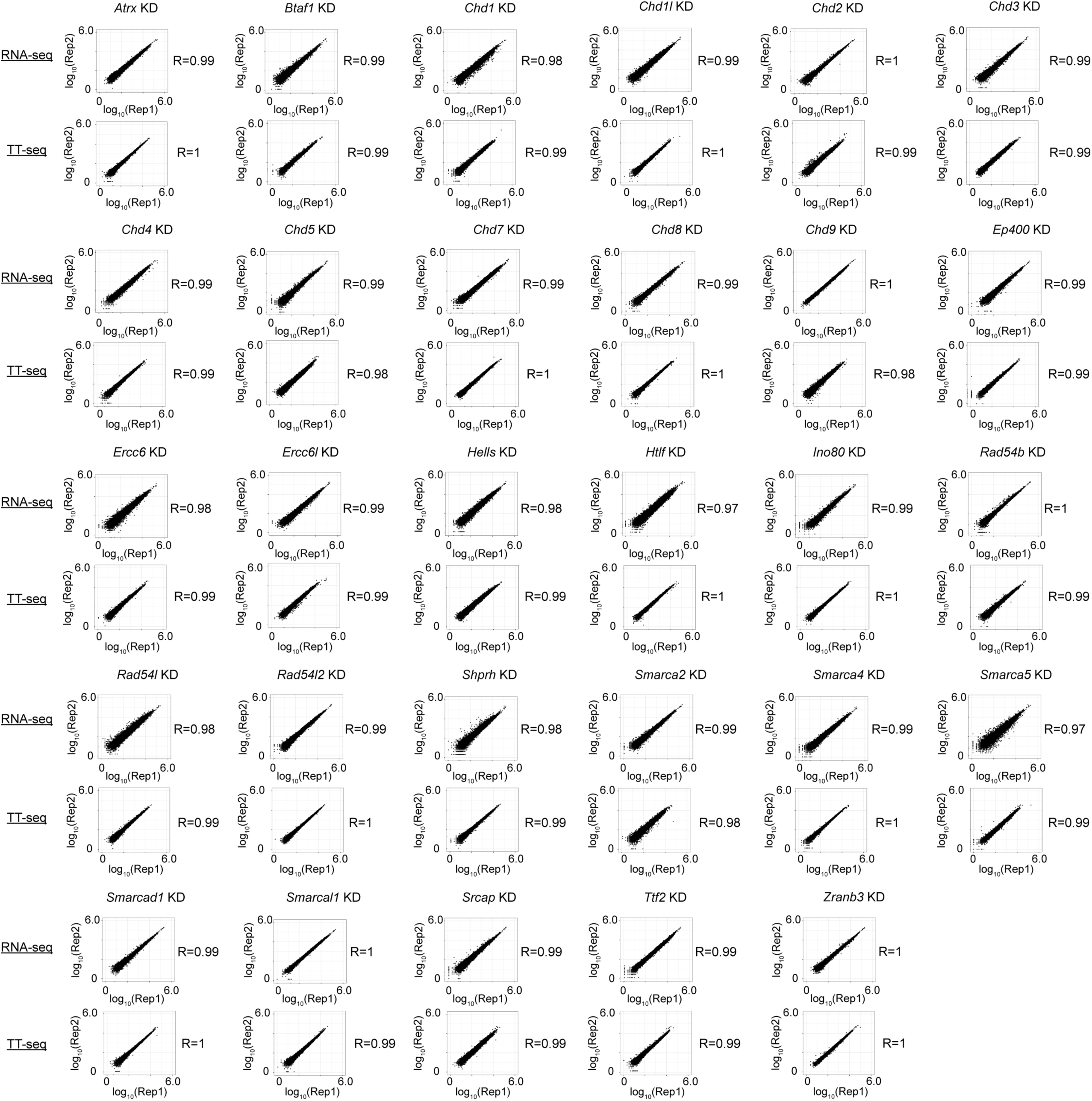
Scatterplots demonstrating high reproducibility of TT-seq and RNA-seq replicates. Related to Figures 1-4. Scatterplots showing correlation of log_10_(DEseq-normalized counts) of Gencode v23 features for biological replicates of each depletion condition in both TT-seq and RNA-seq datasets. Spearman correlation is listed next to the plot.

**Figure S2:**
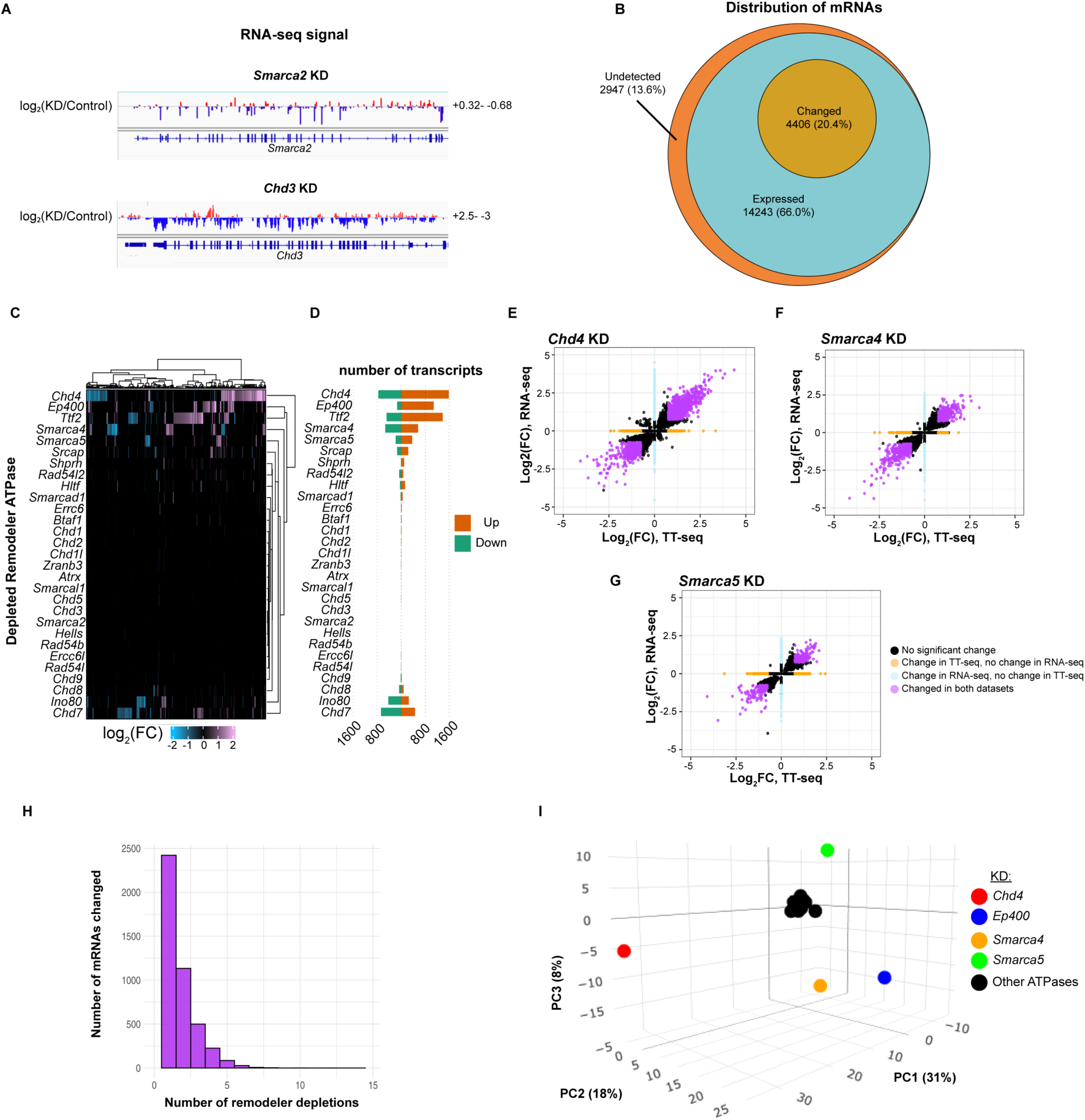
Remodelers alter mRNA transcription throughout the ES cell genome. Related to Figure 1. A. Browser tracks of replicate averaged RNA-seq showing the log_2_(KD/control) over the target gene body. *Smarca2* depletion is shown in the upper panel and *Chd3* depletion is shown in the bottom panel. B. Venn diagram showing the distribution of mRNAs either undetected, expressed, or with changed transcription in this study, identified across all TT-seq datasets. C. Heatmap showing the change in mRNA expression for genes changed across all 29 RNA-seq datasets (|log_2_(FC)|≥0.75 and FDR≤0.05). n=5,810 transcripts. D. Barplot quantifying the number of mRNAs upregulated (orange) or downregulated (green) upon remodeler depletion using RNA-seq (|log_2_(FC)|≥0.75 and FDR≤0.05). E. Scatterplot showing distribution of mRNAs with significant changes in transcription (TT-seq) and/or RNA-seq datasets for *Chd4* depletion (|log_2_(FC)|≥0.75 and FDR≤0.05). Purple dots represent mRNAs with correlated changes in both datasets, orange dots represent mRNAs with significant changes in TT-seq data only, blue dots represent mRNAs with significant changes in RNA-seq data only, and black dots represent mRNAs with not significant change in either dataset. F. As in D, for *Smarca4* depletion. G. As in D, for *Smarca5* depletion. H. Barplot showing the distribution of mRNAs with altered transcription in one or more depletion dataset using TT-seq (|log_2_(FC)|≥0.75 and FDR≤0.05). I. PCA plot of differential transcription data from TT-seq where mRNAs changed in 3 or more depletions. n=851 transcripts.

**Figure S3:**
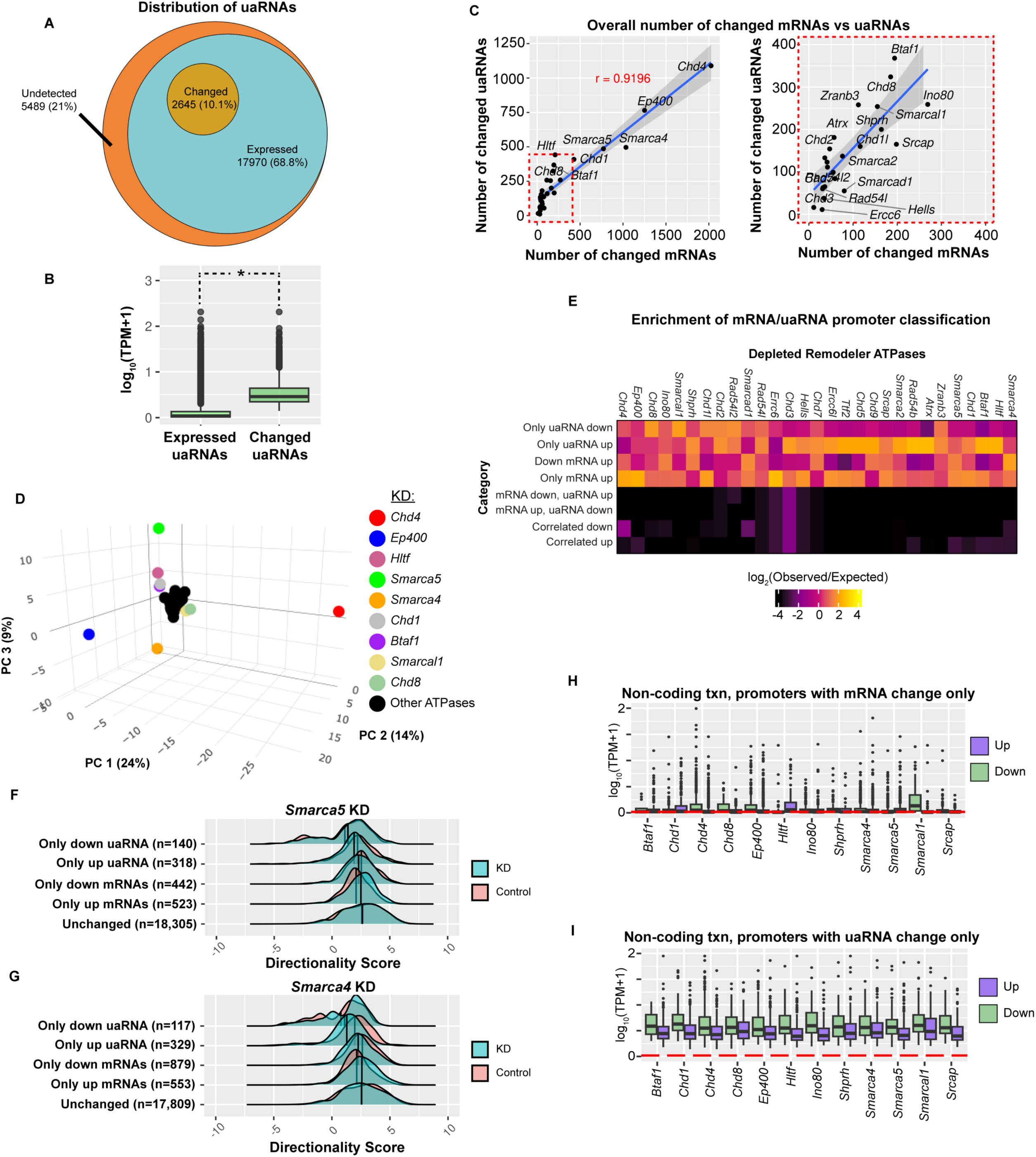
uaRNA transcription is regulated by many SNF2-type remodelers. Related to Figure 2 and 3. A. Venn diagram showing the distribution of uaRNAs either undetected, expressed, or with changed transcription in this study, identified across all TT-seq datasets. B. Boxplots comparing the relative transcription levels of uaRNAs with observable levels of transcription and significant changes in transcription (log_2_(FC)≥0.5 and FDR≤0.05) in any depletion TT-seq dataset. The black line represents the median and edges represent the first and third quartiles. C. Scatterplot comparing the numbers of mRNAs and uaRNAs with significant change in each depletion condition quantified using TT-seq. The right panel is a zoom in containing the region outlined in the dotted red box in the left panel. mRNA change |log_2_(FC)|≥0.75 and FDR≤0.05 and uaRNA change |log2(FC)|≥0.5 and FDR≤0.05. D. PCA plot of differential transcription data from TT-seq where uaRNAs are changed in 3 or more depletions. n=1,118 transcripts. E. Heatmap showing the ratio of observed over expected enrichment of promoters sorted into eight categories based on mRNA and uaRNA change in all depletions, tested by Chi-squared test for homogeneity. Yellow represents higher enrichment and black represents lower enrichment than expected. F. Ridge plot comparing the directionality scores of promoters with significant changes in mRNA and/or uaRNA transcription from TT-seq in *Smarca5* depletion (blue) relative to control (pink). The black line represents the median value. G. As in E, for *Smarca4* depletion. H. Box plot quantifying wildtype ES cell uaRNA expression (TT-seq) at promoters showing only up (purple) or down (green) mRNA transcription in the indicated depletion. I. As in J, for promoters showing only up or down uaRNA expression.

**Figure S4:**
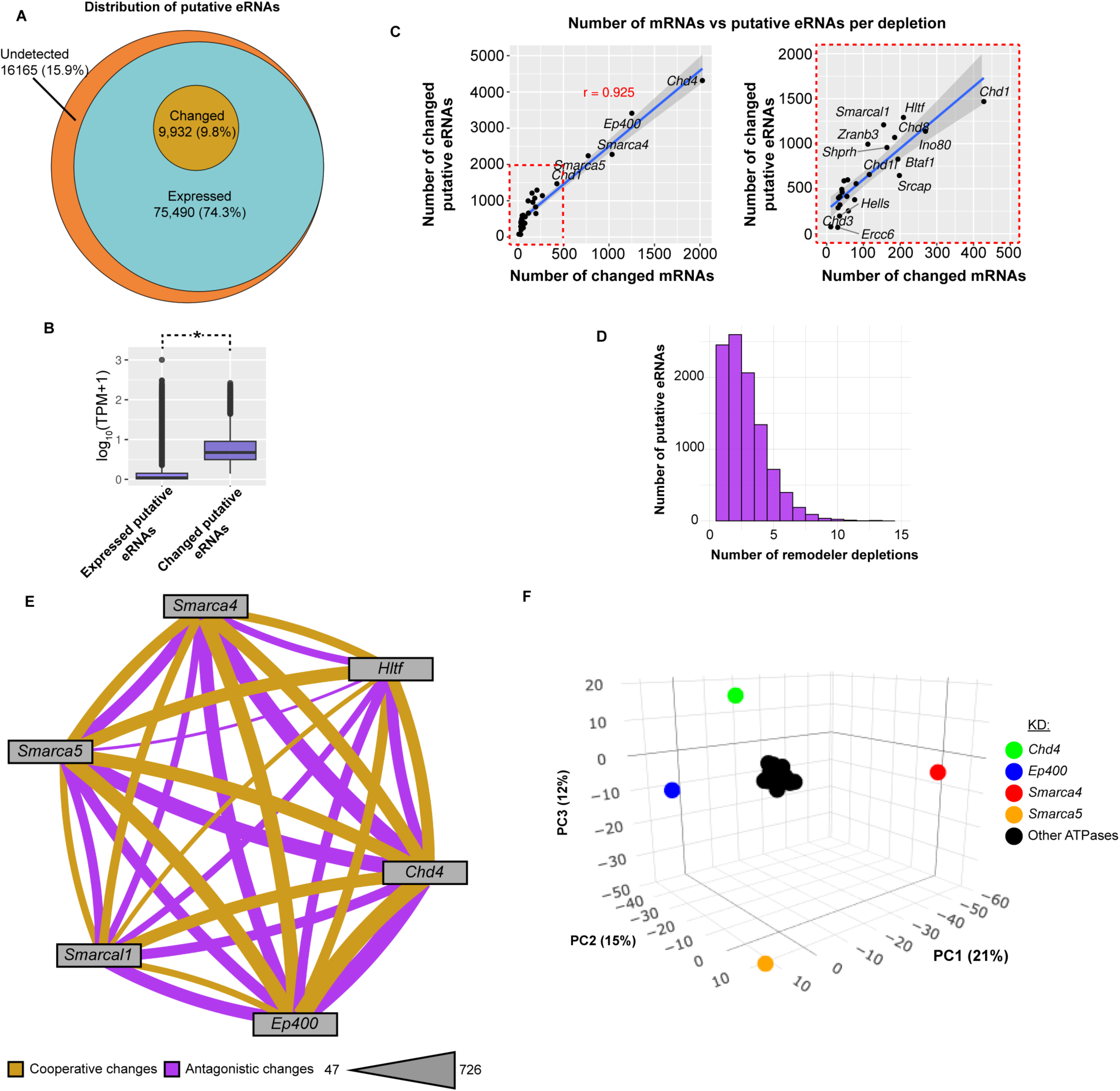
Many remodelers influence eRNA transcription in ES cells. Related to Figure 4. A. Venn diagram showing the distribution of putative eRNAs either undetected, expressed, or with changed transcription in this study, identified across all TT-seq datasets. B. Boxplots comparing the relative transcription levels of putative eRNAs with observable levels of transcription and significant changes in transcription (log_2_(FC)≥0.5 and FDR≤0.05) in any depletion TT-seq dataset. The black line represents the median and edges represent the first and third quartiles. C. Scatterplot comparing the numbers of mRNAs and putative eRNAs with significant changes in each depletion condition quantified using TT-seq. The right panel is a zoom in containing the region outlined in the dotted red box from the left panel. mRNA change |log_2_(FC)|≥0.75 and FDR≤0.05 and putative enhancer change |log2(FC)|≥0.5 and FDR≤0.05. D. Barplot showing the distribution of putative eRNAs with altered transcription (|log_2_(FC)|≥0.75 and FDR≤0.05) in one or more depletion datasets quantified using TT-seq. E. Network representing the number of putative eRNAs with altered transcription (|log_2_(FC)|≥0.5 and FDR≤0.05) in the same direction (cooperative, purple) or opposite direction (antagonistic, yellow) shared between six depletion datasets. Thickness of line correlates with number of putative enhancers in category shared between datasets, with the range listed. F. PCA plot of differential transcription data from TT-seq where putative eRNAs are changed in 3 or more depletions. N=4,882 transcripts.

**Figure S5:**
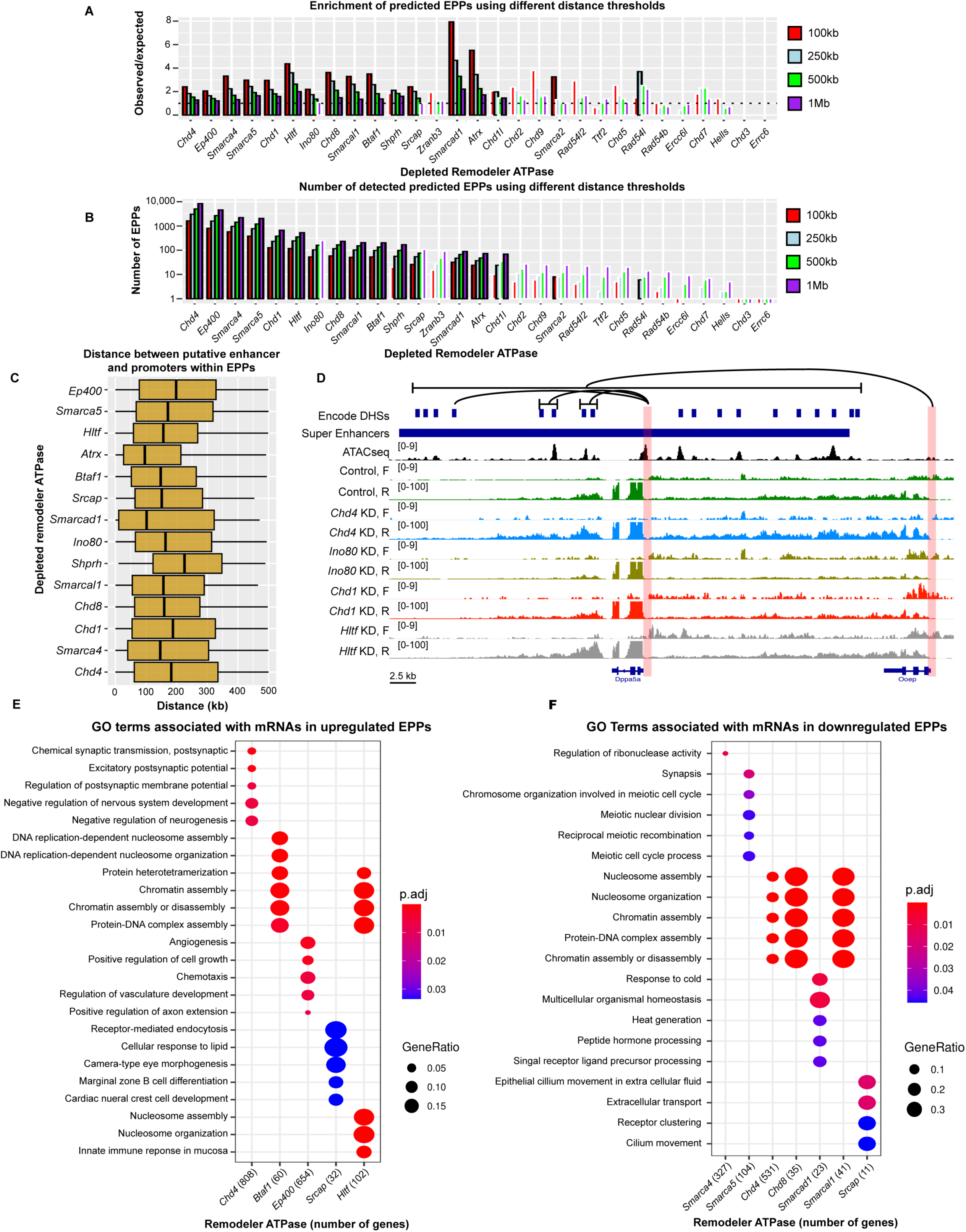
Characterization of EPPs detected in 14 remodeler depletions. Related to Figure 5. A. Barplot showing the ratio of observed predicted enhancer-promoter pairs (EPPs) in each depletion using different distance thresholds over expected numbers determined by permutation test (n=1000). Bars with black outlines represent observed numbers significantly different from expected numbers determined by permutation test (Adjusted *P* value≤0.05). TT-seq. B. Barplot showing the number of predicted EPPs in each depletion quantified with TT-seq using different distance thresholds. Bars with black outlines represent observed numbers significantly different from expected numbers determined by permutation test (Adjusted *P* value≤0.05). C. Boxplots representing the distribution of distances between predicted EPPs in each depletion using a 500 kB distance threshold. The black line represents the median and edges represent the first and third quartiles. D. Browser track showing chromatin accessibility (ATACseq) and comparing transcription (TT-seq) in the control, *Chd4* KD, *Ino80* KD, *Chd1* KD, and *Hltf* KD over the *Dppa5a* locus, *Ooep* locus, and the interacting nearby super enhancer region. Red boxes indicate promoter regions of each locus, and lines indicate DHSs within the super enhancer region shown to interact with each promoter. E. GO terms associated with mRNAs in predicted upregulated EPPs in *Chd4*, *Btaf1*, *Ep400*, *Srcap*, and *Hltf* depletions quantified using TT-seq. Number in parenthesis indicates the number of mRNAs in predicted upregulated EPPs in that depletion. Size of dot indicates relative proportion of total mRNAs associated with indicated GO term and color indicates significance of the association. F. As in D, for GO terms associated with mRNAs in predicted downregulated EPPs in *Smarca4*, *Smarca5*, *Chd4*, *Chd8*, *Smarcad1*, *Smarcal1*, and *Srcap* depletions.

**Figure S6:**
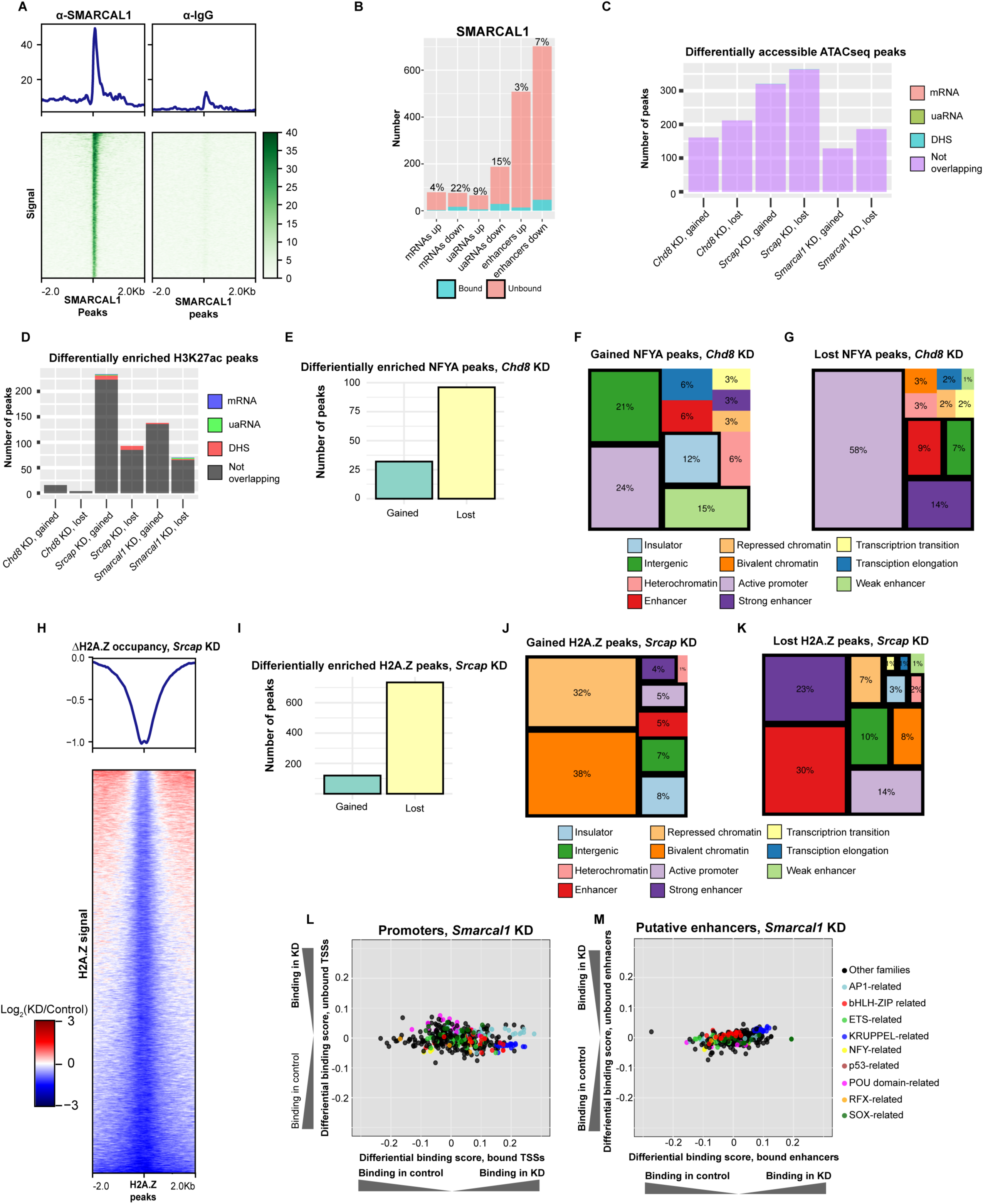
Characterization of chromatin changes defined upon depletion of *Srcap*, *Chd8*, or *Smarcal1* in ES cells. Related to Figure 5 and 6. A. Heatmap showing SMARCAL1 binding over called peaks relative to IgG using CUT&RUN. n= 2,271 peaks. B. Barplot showing the number of CREs within each altered transcript class bound by SMARCAL1 using CUT&RUN. Blue represents number of bound elements and pink indicates number of unbound elements. C. Barplot showing the number of ATAC-seq peaks with differential accessibility (|log_2_(FC)|≥0.5 and FDR≤0.05) in *Chd8*, *Srcap*, or *Smarcal1* depletions, and the number of peaks that overlap promoters or enhancers with correlated significant changes in mRNA, uaRNA, or putative eRNA transcription quantified with TT-seq. D. Barplot showing the number of H3K27ac CUT&RUN peaks with differential enrichment (|log_2_(FC)|≥0.5 and FDR≤0.05) in *Chd8*, *Srcap*, or *Smarcal1* depletions, and the number of peaks that overlap promoters or enhancers with correlated significant changes in mRNA, uaRNA, or putative eRNA transcription quantified with TT-seq. E. Barplot showing the number of gained or lost differentially enriched NFYA CUT&RUN peaks in *Chd8* depletion relative to control. F. Treemap showing the distribution of chromatin states (defined by ChromHMM^84^) that overlap with NFYA gained CUT&RUN peaks in the *Chd8* depletion. Relative size and percentage represent the fraction of total peaks overlapping each chromatin state. States outlined in black represent significantly enriched categories (Fisher’s test, Bonferroni adjusted P value ≤0.05). G. As in D, for NFYA CUT&RUN peaks lost upon *Chd8* depletion. H. Heatmap showing the change in H2A.Z binding over called H2A.Z peaks in the *Srcap* KD relative to control CUT&RUN experiments. n=29,950 peaks. I. As in C, for gained and lost H2A.Z CUT&RUN peaks upon *Srcap* depletion. J. As in D, for H2A.Z CUT&RUN peaks gained upon *Srcap* depletion. K. As in D, for H2A.Z CUT&RUN peaks lost upon *Srcap* depletion. L. Scatterplot comparing the binding score of different transcription factor motifs at promoters bound versus not bound by CHD8. Motifs representing factors from similar related groups are colored according to the legend. All factors shown have *P*≤0.05 at bound and unbound loci, defined from ATAC-seq data using TOBIAS. M. As in L, for putative enhancers bound by SMARCAL1.

## Supplementary information

- Table S1: RT-qPCR and DEseq2 remodeler results
- Table S2: Table containing overall numbers changed under each remodeler
- Table S3: Table contains overlap numbers of overlap for each transcript class
- Table S4: Oligos datasheet esiRNA target sequences and primers
- Table S5: Sample sequencing information chart
- Table S6: Combat-seq batch information

## Acknowledgements

We thank Tom Fazzio and members of the Hainer Lab, especially Dave Klein, for critical comments, discussion, and feedback regarding this article. This work was supported by the NIH grant R35GM133732 (to S.J. Hainer) and NSF GRFP (to B. J. Patty). This project used the Illumina NextSeq2000 available at the University of Pittsburgh Health Sciences Sequencing Core at UPMC Children’s Hospital of Pittsburgh for sequencing, with special thanks to its director, William MacDonald. This research was supported in part by the resources provided by the University of Pittsburgh Center for Research Computing.

## Key resources table

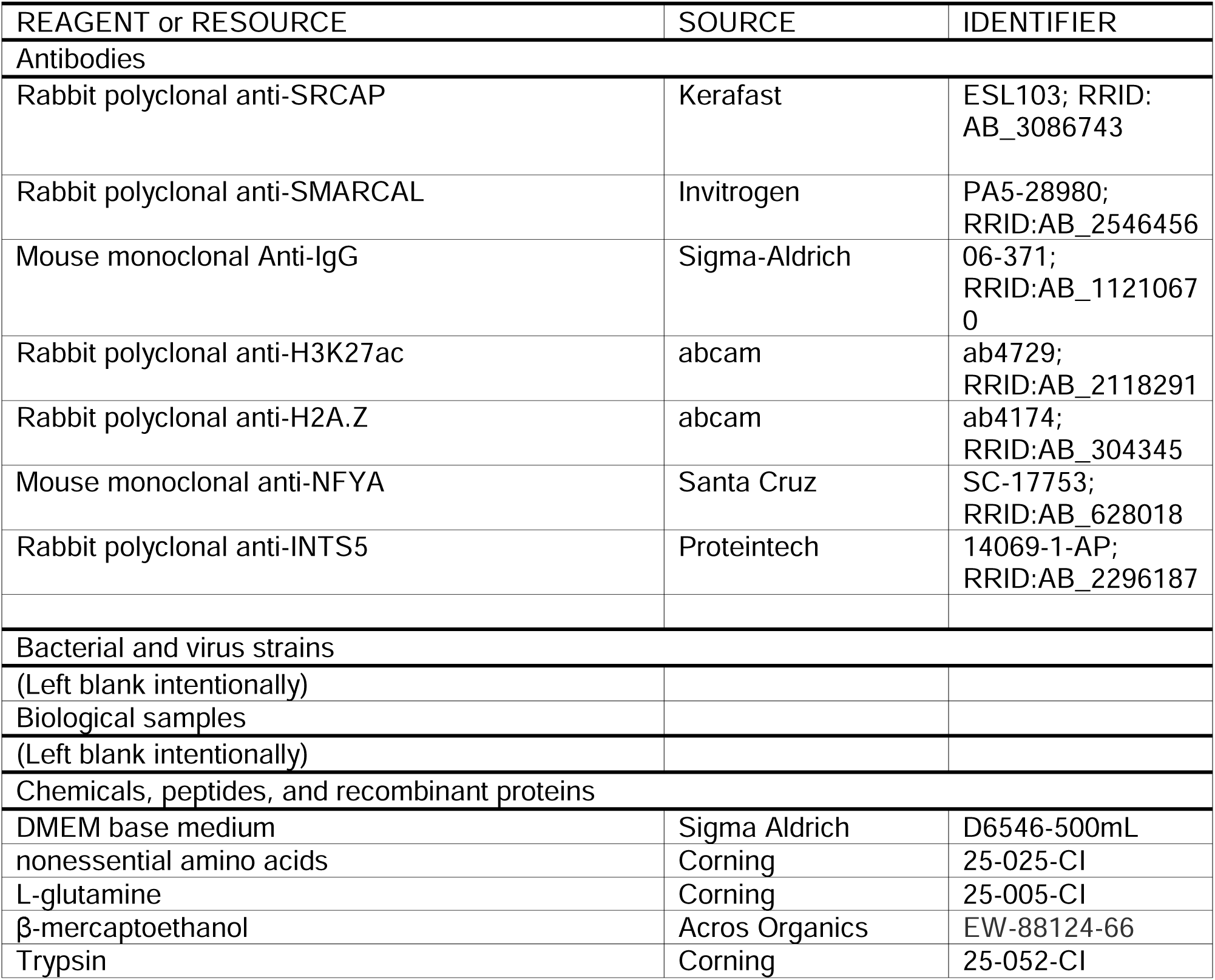
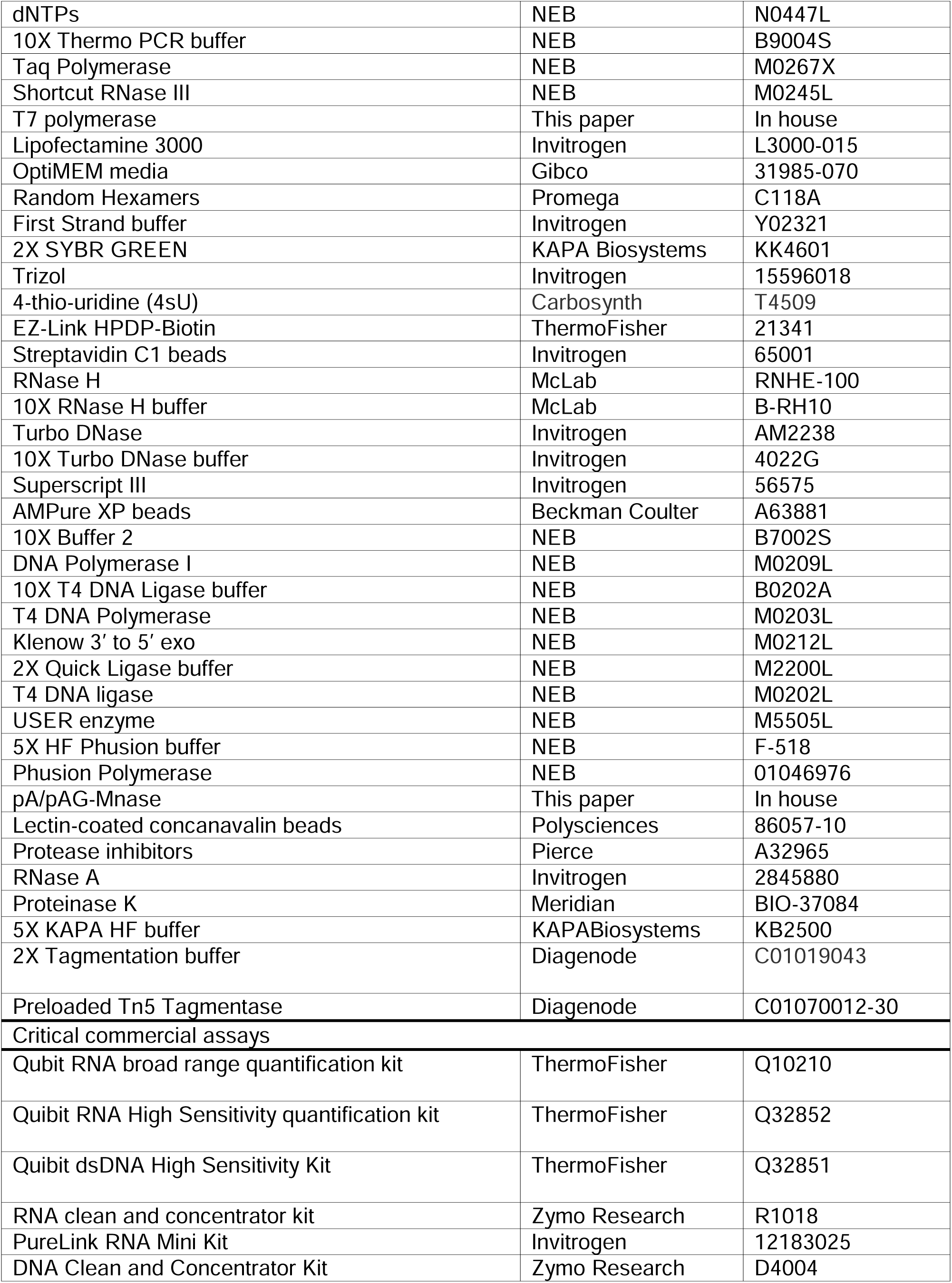
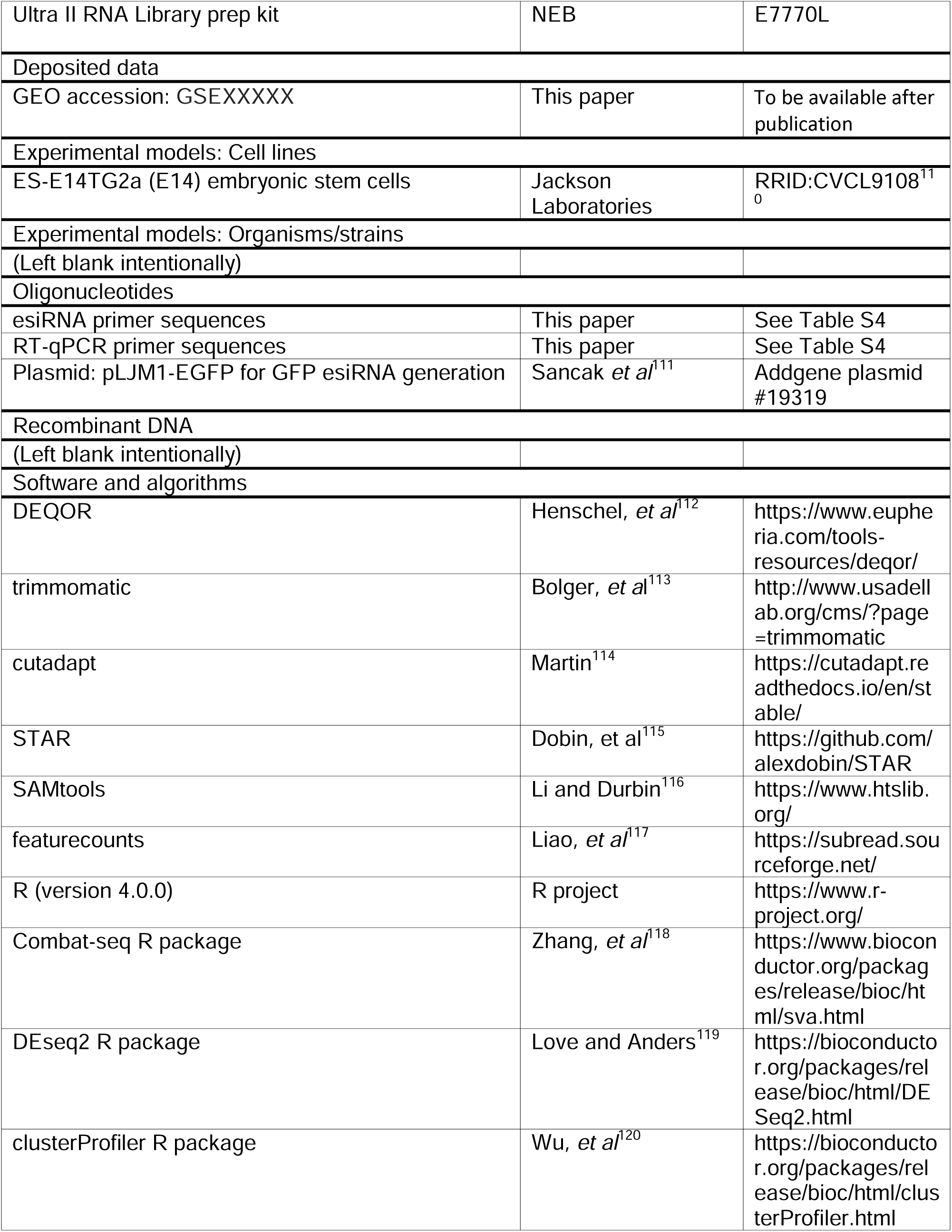
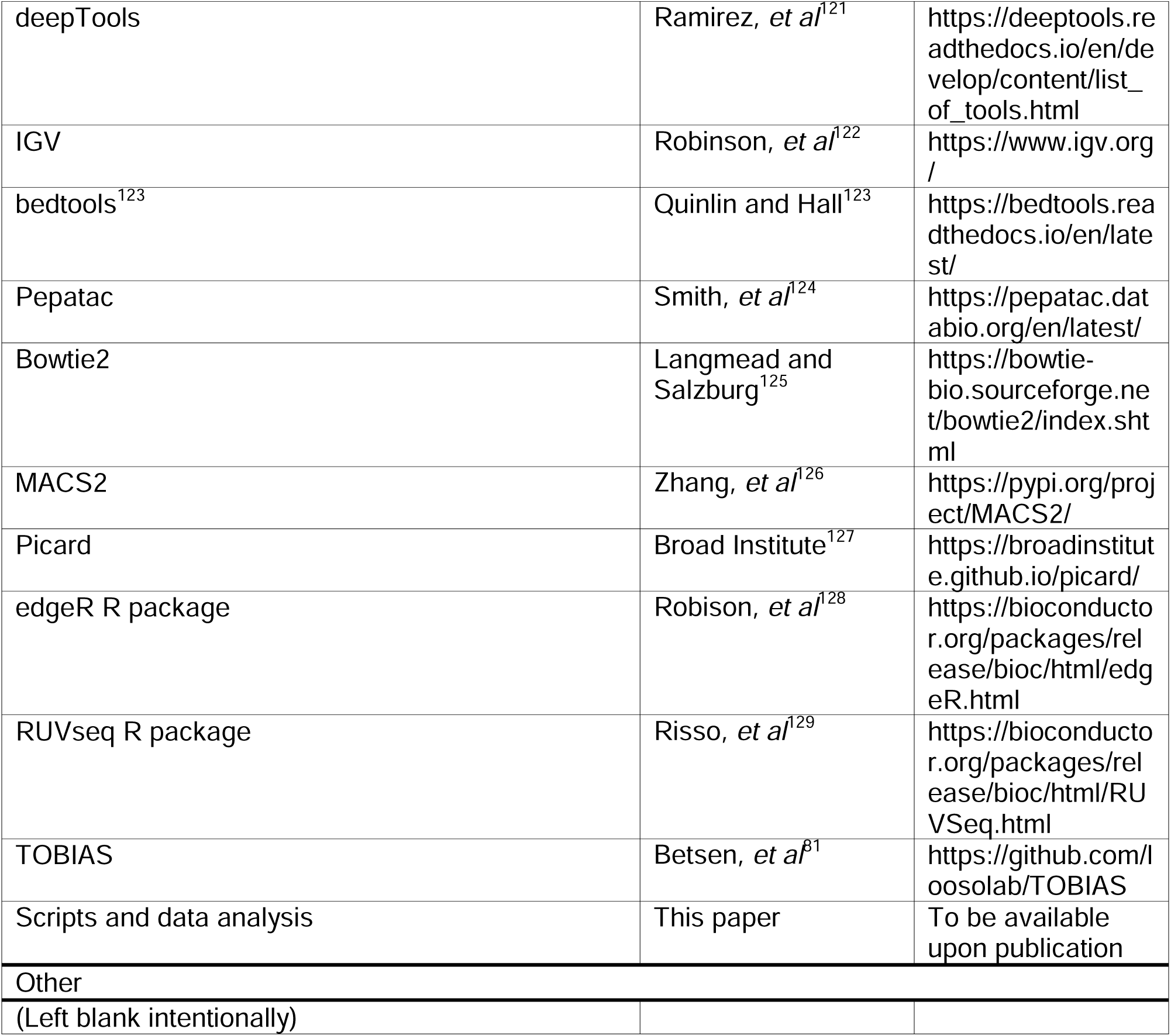

## RESOURCE AVAILIBILITY

### Lead contact

All information and resource request for resources should be directed to the lead contact, Sarah Hainer.

### Materials availability

All unique materials and resources generated from this study are available upon request from the lead contact.

### Data and code availability

- Raw data and processed files have been deposited on GEO accession and will be publicly available at time of publication.
- All original code and data analyses have deposited on Github and will publicly available at time of publication. DOIs are listed in the key resources table.
- Any further needed information is available upon request from the lead contact.

## EXPERIMENTAL MODEL AND SUBJECT DETAILS

### Cell culture

ES-E14TG2a (E14) embryonic stem cells from male *Mus musculus* origin (RRID:CVCL9108^110^) were grown at 37°C and 5% CO_2_ in DMEM base medium supplemented with 10% FBS, 1X nonessential amino acids, 2mM L-glutamine, β-mercaptoethanol and LIF on 10-cm plates precoated with 0.2% gelatin. Cells were passaged every ∼48 h using trypsin and split at a ratio of ∼1:8 with fresh medium. Routine anti-mycoplasma TC hood cleaning was conducted (LookOut DNA Erase spray) and cell lines were screened to confirm no mycoplasma presence.

## METHOD DETAILS

### esiRNA generation

Endoribonuclease-digested short interfering RNAs (esiRNAs) were generated as previously described^14,130^. esiRNAs for all genes targeted in this study were designed and produced using the following protocol. Sequences for esiRNA design were selected by downloading and identifying common exons shared by all confirmed transcript isoforms of each remodeler ATPase according to NCBI. Next, these regions were used as input into DEQOR^112^ and regions predicted to have no off-targets and high likelihood of knockdown efficiency were selected (Table S4). Using the identified regions, oligos were designed containing the T7 promoter sequence (Table S4). Next, the following primary PCR reaction was assembled in 200 µL tubes: 14 µL of nuclease-free water, 0.5 µL of 10 µM target specific forward primer, 0.5 µL of 10 µM target specific reverse primer, 0.5 µL of 10 mM dNTPs, 2 µL of 10X Thermo PCR buffer (NEB), 2 µL of ES-E14 wildtype cDNA template or from 1 µL plasmid pLJM1-EGFP for GFP esiRNAs, and 0.5 µL of Taq Polymerase (NEB), and the following PCR reaction was performed: 1) 94°C for 5 min, 2) 94°C for 30 seconds, 3) 65°C for 30 seconds, 4) 72°C for 2 min, repeating steps 2-4 for 35 total cycles, then 72°C for 2 min. The primary PCR product was diluted 1:200 in nuclease-free water and used as a template for the secondary PCR reaction as follows: 40.5 µL of nuclease free water, 5 µL of 10X Thermo PCR buffer, 1 µL of 10 mM dNTPs, 2 µL of 10 mM T7 primer, 0.5 µL of Taq Polymerase, 1 µL of 1:200 primary PCR product. Secondary PCR reactions were amplified using the following parameters: 1) 94°C for 2 min, 2) 94°C for 30 seconds, 3) 42° for 45 seconds, 4) 72°C for 1 min, repeating steps 2-4 for 5 total cycles, 5) 94°C for 30 seconds, 6) 60°C for 45 seconds, and 7) 72°C for 1 min, repeating steps 5-7 for 30 total cycles, then 7) 72°C for 5 min. Secondary PCR reactions were combined, ethanol/salt precipitated, and resuspended in 50 µL nuclease-free water. Combined and precipitated secondary PCR product was then used as template for the following in vitro transcription (IVT) reaction: 9 µL of combined and precipitated secondary PCR product, 6 µL of 25 mM NTPs, 4 µL of 5X T7 buffer (0.4 M HEPES pH 7.6, 0.2 M DTT, 120 mM MgCl_2_, 10 mM Spermidine), and 1 µL of T7 Polymerase. The IVT reactions were incubated in a thermocycler using the following program: 1) 37°C for 5 h 30 min, 2) 90°C for 3 min, 3) ramp down (0.1°C/sec) to 70°C, 4) 70°C for 3 min, 5) ramp down (0.1°C/sec) to 50°C, 6) 50°C for 3 min, 7) ramp down (0.1°C/sec) to 25°C, and then 8) 25°C for 3 min. Next, IVT products were treated with 1 U of DNase I (NEB) at 37°C for 15 min and brought to a final volume of 100 µL with nuclease-free water. Finally, IVT products were digested with Shortcut RNase III (NEB) and purified using the PureLink RNA Mini Kit (Invitrogen) according to the following modified protocol: 200 µL of Lysis buffer (from RNA Mini Kit) was added to the IVT products, vortexed for 10 seconds, then 260 µL of 100% isopropanol was added and vortexed for an additional 15 seconds. Samples were then applied to a supplied spin column (from RNA Mini Kit), centrifuged for 30 seconds at 10,000 rcf, and the flowthrough was transferred to a new 1.5 mL tube, discarding the column. 700 µL of 100% isopropanol was added to the flowthrough, and vortexed for 15 seconds. 700 µL of the sample was applied to a new spin column, centrifuged for 30 seconds at 10,000 rcf, and the flowthrough was discarded. This process was repeated until the entire sample was applied to the column. 500 µL of Wash buffer 2 (from RNA Mini Kit) was added to the column, which was centrifuged for 30 seconds at 10,000 rcf. Then, 30 µL of nuclease-free water was applied to column, and samples were incubated for 1 min at room temperature, then centrifuged for 30 seconds at 10,000 rcf. This process was repeated a second time for a total eluent volume of 60 µL. Final esiRNAs were quantified using a Nanodrop and visualized on a 1.5% agarose gel to assure no undigested product remained.

### esiRNA transfection

Reverse transfections were performed using 3500-5000 ng of either target or control (GFP) esiRNAs, 25 µL of Lipofectamine 3000, and 2 mL of OptiMEM media, incubated for 15-30 min at room temperature (RT). During this incubation, ES cells were counted and diluted to 350,000 cells/mL in ES cell media. After incubation, the transfection reaction was added to 4 mL diluted cell suspension and transferred to a pre-gelatinized 10 cm plate. After 16-18 h, the media was replaced with 6 mL fresh ES cell media. Cells were either treated with 4sU or harvested 48 h post-transfection for downstream assays, described below.

### RT-qPCR

RNA isolated from transfected cells was quantified using a Nanodrop, and 1 μg of RNA was mixed with 2 μL of 10 mM dNTP mixture, 1 μL of random hexamers (Promega), and brought to 20 μL total volume with nuclease-free water. Samples were then incubated in a thermocycler at 68°C for 5 min, then placed on ice for 2 min. Then, 8 μL of First Strand buffer (Invitrogen), 4 μL of 0.1M DTT, 7 μL of nuclease-free water, and 1 μL of homemade reverse transcriptase (RT) were added to each sample, mixed by pipetting, and samples were placed into a thermocycler. The following program was used: 42°C for 90 min and 70°C for 15 min. For each sample, three technical replicates of 1 μL cDNA, 5 μL 2x SYBR GREEN, 2 μL nuclease-free water, and 1 μL 5 mM sample-specific forward and reverse qPCR primers (Table S4) were combined and run on a Roche Light Cycler for 25 cycles. Abundance of the target transcript in depleted samples was determined using the ΔΔCT normalization method relative to control samples, using *Gapdh* transcript abundance for internal normalization as previously described^131^.

### Transient Transcriptome sequencing (TT-seq)

TT-seq was conducted as previously described^132,133^. 48 h post transfection, media was aspirated from transfected plates and replaced with 10 mL of 500 nM 4-thio-uridine (4sU) containing ESC media and the plates incubated at 37°C with 5% CO_2_ for 5 min. After 5 min, the 4sU-containing media was aspirated and the cells were washed with PBS, typsinized, and then pelleted by centrifugation. Total RNA was collected from cell pellets with a TRIzol extraction followed by an isopropanol/salt precipitation and resuspended in 100 µL 1XTE, according to ThermoFisher’s recommendations. RNA concentration was determined by Qubit with the Qubit RNA broad range quantification kit (ThermoFisher). 1 µg of RNA was used as a template for RT-qPCR to determine esiRNA depletion efficiency, as described above. 100 µg of total RNA was diluted to a concentration of 240 ng/µL at a volume of 416.67 µL in 1XTE and then fragmented with a Bioruptor Pico (Diagenode) on high power for one 30 second cycle. The fragmented RNA was then combined with 283.33 µL 1XTE, 100 µL 10X Biotinylation buffer (100 mM Tris pH 7.4 and 10 mM EDTA), and 200 µL of 1 mg/mL biotin-HPDP (ThermoFisher) in dimethylformamide (DMF; freshly prepared). Samples were vortexed, then incubated in a thermomixer at 37°C shaking at 1000 RPM in the dark for 2 h. Samples were then chloroform extracted, isopropanol/salt precipitated, and resuspended in 22 µL of nuclease-free water. Streptavidin C1 beads (Invitrogen) were prepared for RNA separation as follows: 60 µL of beads were rotated for 2 min at room temperature RT with 1 mL of 1 M NaOH and 50 mM NaCl and then placed in the magnetic rack for 1 min. The supernatant was discarded, and beads were resuspended in 1 mL of 100 mM NaCl. Beads were washed twice with 1 mL of 100 mM NaCl and resuspended in 60 µL of TT-seq Binding buffer (10 nM Tris pH 7.4, 300 mM NaCl, 0.1% Triton). Then, 60 µL of prepared streptavidin C1 beads were added to each sample and rotated at room temperature for 20 min. Following incubation, the samples were magnetized for 1 min and the supernatant (containing the unlabeled RNA) was placed in a separate 1.5 mL tube and put on ice. The unlabeled RNA from supernatant was Phenol:Chroloform:Isopropanol(PCI)/chloroform extracted, isopropanol/salt precipitated, and resuspended in 100 µL of nuclease-free water. The bead-bound labeled nascent RNA was washed twice with 500 µL of High Salt buffer (50 mM Tris pH 7.4, 2M NaCl, 0.5% Triton), twice with 500 µL of TT-seq Binding buffer, and once with 500 µL of Low Salt buffer (5 mM Tris pH 7.4, 0.1% Triton), rotating for 1 min at RT, re-magnetizing and resuspending the beads during each wash. The nascent RNA was eluted by resuspending beads in 100 µL of freshly prepared 100 mM DTT and incubating in a thermomixer at 65°C and 1000 RPM shaking for 5 min. The beads were magnetized and the supernatant was transferred to a new tube, and a second elution from the beads was performed. Eluted nascent RNA were pooled and the nascent RNA was recovered with a PCI extraction and an isopropanol/salt/glycogen precipitation. RNA pellets were resuspended in 25 µL of nuclease-free water. The total RNA and nascent RNA from each sample were used to build RNA-seq and TT-seq libraries, respectively, as described below.

### RNA-seq library preparation

RNA-seq libraries were built using a custom strand-specific RNA-seq library build protocol that includes an antisense oligo (ASO)-based rRNA depletion protocol. First, 2 µg of total RNA was combined with 2 µL of 0.5 µM pooled antisense rRNA oligos (at a ratio of 1 µL of 0.5 µM pooled antisense rRNA oligos per µg total RNA) and rRNA hybridization buffer (100 mM Tris-Cl pH 7.4, 200 mM NaCl) to a final volume of 10 µL. In a thermocycler, the samples were heated at 95°C for 5 min, then slowly cooled down to 22°C at a rate of −0.1°C/sec, followed by incubation at 22°C for 5 min, then placed on ice. Next, 2 µL of thermostable RNase H (10 units; Epicentre), 2 µL of 10X RNase H buffer (Epicentre) and 6 µL of nuclease-free water were added and the samples were incubated at 45°C for 30 min. Then, 2 µL of Turbo DNase (ThermoFisher) and 5 µL of 10X Turbo DNase buffer (ThermoFisher) were added and samples were incubated at 37°C for 20 min. Samples were then purified using an RNA clean and concentrator kit (Zymo Research), according to the following protocol. Adjusted RNA Binding buffer was made by combining 50 µL of RNA Binding buffer with 50 µL of 100% ethanol and mixing well by pipetting or vortexing. Then, 100 µL of adjusted RNA Binding buffer was added to each sample and mixed well by vortexing, and then each sample was applied to a spin column supplied in the RNA clean and concentrator kit (Zymo Research). Samples were centrifuged at 12,500 rcf for 30 seconds, and the flowthrough was discarded. 400 µL of RNA Prep buffer was added and samples were centrifuged at 12,500 rcf for 30 seconds, and the flowthrough was discarded. 700 µL of RNA Wash buffer was added and samples were centrifuged at 12,500 rcf for 30 seconds, and the flowthrough was discarded. 400 µL of RNA Wash buffer was added and samples were centrifuged at 12,500 rcf for 2 min, and flowthrough was discarded. Samples were centrifuged again at 12,500 rcf for 1 min to remove any residual RNA Wash buffer, and the columns were transferred to fresh 1.5 mL microfuge tubes. 11 µL of nuclease-free water was added to each column and incubated for 1 min at room temperature before spinning at 12,500 rcf for 1 min. Samples were transferred to 200 µL tubes, 5 µL 5X First Strand buffer (Invitrogen) was added, and the samples were heated at 95°C for 5 min in a thermocycler. Then, 1 µL of 6 M random hexamers were added to each sample and incubated at 65°C for 3 min. Next, 5.25 μL of nuclease-free water, 1.5 μL of 10 mM dNTPs, 1.25 μL of 100 mM DTT, and 1 μL of Superscript III (Invitrogen) were added to each sample, and then placed in a thermocycler with the following program: 25°C for 5 min, 50°C for 1 h, and 70°C for 15 min. Next, samples were purified using 45 µL AMPure XP beads (Beckman Coulter) according to the manufacturer’s instructions and eluted in 22 µL of 0.1X TE. 20 μL of eluted sample was transferred to fresh 200 μL tubes. To each sample, 3 μL of 10X NEB buffer 2, 2 μL of dUTP mixture (20 mM dUTP, 10 mM dATP, 10 mM dCTP, 10 mM dGTP), 0.5 μL of 100 mM DTT, 1 μL of RNase H (Epicentre), and 2 μL of DNA Polymerase I (10U/μL, NEB) were added and mixed by pipetting, then the samples were incubated in a thermocycler at 16°C for 2.5 h. Samples were then purified with 45 µL AMPure XP beads (Beckman Coulter) according to the manufacturer’s instructions and eluted with 32 µL 0.1X TE. 30 μL of eluted sample was transferred to fresh 200 μL tubes. To each sample, 3 μL of nuclease-free water, 5 μL of 10X T4 DNA Ligase buffer (with 10mM ATP), 5 μL of 10 mM dNTP mix, 2 μL of T4 DNA Polymerase (3 U/μL, NEB) 1 μL of Klenow DNA Polymerase (5 U/μL, NEB) and 2 μL of T4 PNK (10 U/μL, NEB) were added, and the samples were incubated in a thermocycler at 20°C for 30 min. Next, 35 µL of AMPure XP beads were added to each sample and mixed by vortex, and incubated at RT for 5 min, then magnetized on a magnetic rack for 3 min. The supernatant was transferred to new 200 µL tubes, and then the samples were purified with 102 µL AMPure XP beads according to the manufacturer’s instructions and eluted with 22 µL 0.1XTE and 20 μL of eluted sample was transferred to fresh 200 μL tubes. To each sample, 9 μL of nuclease-free water, 5 μL of NEB buffer 2, 1 μL of 10 mM dATP, and 3 μL of Klenow 3’ to 5’ exo (5 U/μL, NEB), and samples were incubated in a thermocycler at 37°C for 30 min. This was followed with a purification with 60 µL AMPure XP beads according to the manufacturer’s instructions, eluted with 22 µL 0.1XTE and transferred to fresh 200 μL tubes. For adapter ligation, NEBNext adapters were thawed on ice and diluted 1:5 in nuclease-free water to final concentration of 5 mM, and then 25 μL of 2X Quick Ligase buffer, 1 μL 10 mM NEBNext adapter, 2 μL of T4 DNA ligase (NEB) were added and incubated at RT for 30 min. 3 µL USER enzyme (NEB) were added to each sample and incubated at 37°C for 15 min and followed by a purification with 50 µL AMPure XP beads according to the manufacturer’s instructions, eluted with 30 µL 0.1XTE and transferred to fresh 200 μL tubes. To each sample, 1 μL of 10 μM NEBNext i7 Primer (Universal), 1 μL of 10 μM NEBNext i5 Primer, 10 μL nuclease-free water, 12 μL of 5X HF Phusion buffer, 2 μL of 10mM dNTPs mixture, and 1 μL of Phusion Polymerase (2 U/μL, NEB) were added and mixed by pipette. Samples were then placed in a thermocycler and the following PCR program was used: 1) 98°C for 30 sec, 2) 98°C for 10 sec, 3) 65°C for 30 sec, 4) 72°C for 30 sec, repeat steps 2-4 for a total of 10 cycles, and then 72°C for 3 min. A final bead cleanup was performed with 54 µL AMPure XP beads according to the manufacturer’s instructions, eluted with 22 µL of 0.1XTE, and 20 uL of elutant was transferred to fresh 1.5 mL microfuge tubes. Libraries were quantified by Qubit with the dsDNA High Sensitivity Kit (ThermoFisher) and run on a Fragment Analyzer to confirm high quality of each library prior to sequencing. Libraries were paired-end sequenced on the Illumina NextSeq 500 platform with standard protocols at the UPMC Children’s Hospital of Pittsburgh to a depth of ∼40,000,000 uniquely mapped reads per sample.

### TT-seq library preparation

TT-seq libraries were built using the NEBNext Ultra II RNA Library prep kit for Illumina (NEB) according to the manufacturer’s recommendations, with specific modifications as described below. 100 ng of nascent RNA was rRNA depleted using antisense oligo rRNA depletion as described for RNA-seq libraries, but with 0.05 µM pooled antisense rRNA oligos to account for the adjusted amount of input material. To account for using rRNA-depleted nascent RNA as input for the library build, we adjusted the volumes of the Fragmentation and Priming mix, First Strand Synthesis Reaction, and Second Strand Synthesis Reaction as follows: Fragmentation and Priming mix: 10 µL of rRNA-Depleted nascent RNA, 8 µL of NEBNext First Strand Synthesis Reaction buffer, 2 µL of Random Primers; First Strand Synthesis Reaction: 20 µL of RNA, 16 µL of NEBNext Strand Specificity Reagent, and 4 µL of NEBNext First Strand Synthesis Enzyme Mix; Second Strand Synthesis Reaction: 40 µL of First Strand Synthesis product, 8 µL of NEBNext Second Strand Synthesis Reaction buffer with 10X dUTP Mix, 4 µL NEBNext Second Strand Synthesis Enzyme Mix, and 28 µL nuclease-free water. RNA was fragmented for 15 min, NEBNext Adaptor and primers were diluted 1:5, and 7 PCR amplification cycles were performed for all TT-seq library builds. Libraries were quantified by Qubit with the dsDNA High Sensitivity kit and run on a Fragment Analyzer to confirm high quality of each library prior to sequencing. TT-seq libraries were paired-end sequenced on the Illumina NextSeq 500 platform with standard protocols at the UPMC Children’s Hospital of Pittsburgh to a depth of ∼40,000,000 uniquely mapped reads per sample.

### CUT&RUN

CUT&RUN experiments were performed as previously described^132,134,135^, with the following modifications. The following antibodies were used in CUT&RUN experiments performed on 100,000 lightly crosslinked wildtype ES cells using a low salt elution method for fragment release: SRCAP (Kerafast ESL103, 1:50) SMARCAL (Invitrogen PA5-28980, lot YF3945097B, 1:50), and IgG (Sigma-Aldrich, 06-371,1:250). The following antibodies were used in low salt elution uncrosslinked CUT&RUN experiments on 100,000 cells transfected by esiRNAs: H3K27ac (abcam ab4729, lot GR3416784-1, 1:100), H2A.Z (abcam ab4174, lot GR3198864-1, 1:100), NFYA (Santa Cruz SC-17753, lot D2522, 1:50) and IgG (Sigma-Aldrich, 06-371,1:250)..

For lightly crosslinked wildtype ES cells: cells were washed with 1X PBS, typsinized, counted, and 1 million cells were collected in 1.5 mL microfuge tubes and placed on ice. Cells were centrifuged at 600 rcf at 4°C for 5 min, and the supernatant was removed without disturbing the pellet. The cell pellet was resuspended gently in 1 mL ESC media + 0.1% formaldehyde, inverted 3 times, and crosslinked for 5 min at RT. The reaction was then quenched with 100 uL of 2.5M glycine. For 48-h post esiRNA transfected cells: cells were washed with 1X PBS, typsinized, counted, and 600,000 cells were collected in 1.5 mL microfuge tubes and placed on ice. Cells were centrifuged at 600 rcf at 4°C for 5 min, and the supernatant was removed without disturbing the pellet. The cell pellet was resuspended gently in 1 mL cold 1X PBS and centrifuged at 600 rcf at 4°C for 5 min, and the supernatant was removed without disturbing the cell pellet. The cells were then gently resuspended in 1 mL cold Nuclear Extraction (NE) buffer (20 mM HEPES-KOH, pH 7.9, 10 mM KCl, 0.5mM spermidine, 0.1% Triton X-100, 20% glycerol, freshly added protease inhibitors) and incubated on ice for 10 min. After incubation, the lysed cells were centrifuged at 600 rcf at 4°C for 5 min, and the supernatant was removed without disturbing the nuclei pellet, and the pellet was flash frozen. Prior to experimentation, pellets were thawed on ice for 5 min and resuspended in 600 μL cold NE buffer.

To prepare lectin-coated concanavalin beads (Polysciences), 25 uL of beads per 100,000 nuclei in each aliquot were combined with 850 μL Binding buffer (20 mM HEPES, pH 7.5, 150 mM NaCl, 0.5 mM spermidine, 0.1% BSA, 2 mM EDTA, fresh protease inhibitors) in a fresh tube, washed twice with 1 mL Binding buffer on a magnetic rack, and resuspended in 300 μL Binding buffer. While gently vortexing the nuclei, 300 μL of bead slurry was slowly added to the cell nuclei and reaction was rotated at 4°C for 10 min. Next, the samples were placed on a magnetic rack until the solution cleared (∼5 min), and the supernatant was removed without disturbing the beads. The beads were resuspended in 1 mL Blocking buffer (20 mM HEPES, pH 7.5, 150 mM NaCl, 0.5 mM spermidine, 0.1% BSA, 2mM EDTA, freshly added protease inhibitors) with gentle pipetting, and then incubated at RT for 5 min. The samples were then placed on the magnet stand, the supernatant was removed and then gently resuspended in 1 mL Wash buffer (20 mM HEPES, pH 7.5, 150 mM NaCl, 0.5 mM spermidine, 0.1% BSA, freshly added protease inhibitors). The samples were placed on a magnetic stand, the supernatant was removed, and resuspended in 125 μL Wash buffer per 100,000 nuclei aliquot and divided into individual 100,000 nuclei aliquots. Next, primary antibody mix (125 μL Wash buffer with target specific antibody at specified dilutions) was added to each sample while gently pipetting and then incubated on a rotator for 1 h at 4°C for lightly crosslinked nuclei or at RT for transfected nuclei. Samples were placed on a magnetic rack, allowed to clear and the supernatant was removed and discarded. Samples were washed twice with 1 mL Wash buffer, resuspending each time by pipetting. After washing, the samples were resuspended in 125 μL Wash buffer, and 125 μL pA-MNase (for rabbit antibodies) or pAG-MNase (for mouse antibodies) mix was added to each sample while gently vortexing. Samples were then rotated for 30 min at 4°C for lightly crosslinked nuclei or at RT for transfected nuclei. Samples were washed twice with 1 mL Wash buffer as above, resuspended in 150 μL Wash buffer and placed in an ice/water bath for 5 min to cool to 0°C. After incubation, 3 μL of 100 mM CaCl_2_ was added to each sample by gently vortexing and flicking 2-3 times to mix well and placed back into the ice bath for 1 h for crosslinked nuclei or 30 min for transfected nuclei. After 1 h or 30 min exactly as indicated, 150 μL of 2xSTOP buffer (200 mM NaCl, 20 mM EDTA, 4 mM EGTA, 1% NP40, 0.2 mg/mL glycogen, and 0.05 ng/mL *S. cerevisiae* DNA spike-in) was added to each sample and mixed by pipetting. Samples were then incubated at 4°C for 1 h to facilitate low-salt fragment release from bead-bound nuclei. Next, samples were placed onto the magnetic rack, and the supernatant was transferred to fresh 1.5 mL microfuge tube without disturbing the beads. Next, 20 μL 5M NaCl and 1.5 μL RNase A (Invitrogen) was added to each sample, mixed well by pipette, then incubated at 37°C for 20 min in a thermomixer. Then, 2.5 μL of Proteinase K and 3 μL of 10% SDS was added, mixed by quick vortex, and then incubated for 10 min at 70°C. At this stage, lightly crosslinked nuclei were then incubated at 55°C overnight to reverse crosslinking. Next, 300 μL of PCI was added to each sample and vortexed for 15 seconds on highest setting. The samples were then transferred to phase lock tubes and centrifuged for 5 min at 4°C at 16,000 rcf. Then, 300 μL of chloroform was added to each sample and vortexed for 15 seconds on the highest setting. The samples were centrifuged for 5 min at 4°C at 16,000 rcf, the aqueous fraction was transferred to a new 1.5 mL microfuge tube. For lightly crosslinked samples, 150 μL of AMPure XP beads were added to the aqueous fraction, mixed well by vortex, and incubated for 15 min at RT. Samples were then magnetized for 5 min at RT, and the supernatant was transferred to a new tube, and 1 mL of 100% ethanol and 5 μL of 20 mg/mL glycogen was added and vortexed for 15 seconds on the highest setting. For transfected nuclei (not crosslinked), the aqueous fraction was transferred to a new tube, and 750 μL of 100% ethanol and 5 μL of 20 mg/mL glycogen were added and vortexed for 15 seconds on the highest setting. The samples were incubated for 30 min at −20°C, and then centrifuged at 16,000 rcf for 30 min at 4°C. The supernatant was discarded, and the DNA pellet was washed with 1 mL of 80% ethanol. Next, the samples were centrifuged at 16,000 rcf for 5 min at 4°C and the supernatant was discarded. Pellets were allowed to air dry for approximately 5 min and resuspended in 50 μL of 0.1XTE.

### CUT&RUN library build

On ice in 200 μL tubes, 7 μL of NEBNext Ultra II End Prep Enzyme buffer (NEB) and 3 μL of NEBNext Ultra II End Prep Enzyme Mix (NEB) were added to each sample and mixed well by pipetting up and down. Samples were placed in a thermocycler with the following program: 20°C for 30 min, then 65°C for 30 min. Next, 5 μL of 1.5 uM NEB adapter, 5 μL of T4 DNA ligase, and 55 μL of 2X Quick Ligase buffer were added to each sample and then mixed by pipetting. Samples were incubated in a thermocycler at 20°C for 15 min, then 3 μL of USER enzyme (NEB) was added and mixed again by pipetting. Samples were incubated in a thermocycler at 37°C for 15 min, then purified with 128 μL of AMPure XP beads (Beckman Coulter) according to the manufacturer’s instructions and eluted in 30 μL of 0.1X TE buffer. After elution, 27.5 μL of each sample were transferred to fresh 200 μL tubes. To each sample, 10 μL of 5X KAPA HF buffer, 1 μL of KAPA HIFI polymerase, 1.5 μL of 10 mM dNTP mix, 5 μL of 1.5 μM NEBNext i5 Universal PCR primer, and 5 μL of 1.5 uM NEBNext i7 PCR primer were added and mixed well by pipetting. Samples were then placed in a thermocycler and then PCR amplified using the following program: 1) 98°C for 45 seconds, 2) 98°C for 15 seconds, 3) 60°C for 10 seconds, repeat steps 2-3 for 15 total cycles, and 4) 72°C for 1 min. Samples were then purified with 60 μL of AMPure XP beads according to the manufacturer’s instructions, eluted in 30 μL of 0.1X TE buffer, and 27.5 μL of amplified library were transferred to fresh tubes. A fraction of each library was run on a 1.5% agarose gel and quantified by Qubit High Sensitivity dsDNA quantification kit to ensure high quality. CUT&RUN libraries were paired-end sequenced on the Illumina NextSeq500 platform with standard protocols at the UPMC Children’s Hospital of Pittsburgh to a depth of ∼10,000,000 uniquely mapped reads per sample.

### ATAC-seq

Cells were transfected with esiRNAs as described above. At 48 h post-transfection, the cells were washed with 1X PBS, typsinized, counted, and 50,000 cells were collected in a 1.5 mL microfuge tube and placed on ice. Cells were centrifuged at 600 rcf at 4°C for 5 min, and the supernatant was removed without disturbing the pellet. The cell pellet was resuspended gently in 1 mL cold 1X PBS, centrifuged at 600 rcf at 4°C for 5 min, and the supernatant was removed without disturbing the cell pellet. The cells were then gently resuspended in 600 μL of cold NE buffer (20 mM HEPES-KOH, pH 7.9, 10 mM KCl, 0.5mM spermidine, 0.1% Triton X-100, 20% glycerol, freshly added protease inhibitors) and incubated on ice for 10 min. After incubation, the samples were centrifuged at 600 rcf at 4°C for 5 min, and the supernatant was removed without disturbing the nuclei pellet. Pellets were flash frozen and stored in the −80°C until use. Prior to use, nuclei were thawed on ice (∼5 min). Nuclei were gently resuspended in 50 μL of Transposase mix (25 μL Transposase buffer, 16.5 μL 1X PBS, 0.5 μL 10% Tween-20, 0.5 μL 1% digitonin, 5 μL nuclease-free water, 2 μL Tagmentase (Diagenode), and then incubated at 37°C for 30 min with 1000 rpm on a thermomixer. After incubation, the samples were isolated using a DNA Clean and Concentrator Kit (Zymo Research) according to the manufacturer’s instructions and eluted in 10 μL of DNA elution buffer.

### ATAC-seq library preparation

To 10 μL of eluted sample in 200 μL tubes, 2.5 μL 25 μM Nextera i7 primer, 2.5 μL 25 μM Nextera i5 primer, 10 μL nuclease-free water, and 25 μL Nextera High Fidelity 2X PCR Master Mix were added and mixed by pipetting. Samples were then PCR amplified using the following program: 1) 72°C for 5 min, 2) 98°C for 30 seconds, 3) 98°C for 10 seconds 4) 63°C 30 seconds, repeat steps 3-4 for 5 total cycles, and 5) 72°C for 1 min. Samples were placed on ice and 1 μL of the partially amplified sample was added to 1 μL 2 μM Nextera primer 1, 1 μL 2 uM Nextera primer 2, 3 μL nuclease-free water, and 5 μL 2X SYBR green and mixed well by pipetting while avoiding air bubbles. qPCR was performed with the following program: 1) 72°C for 5 min, 2) 98°C for 30 seconds, 3) 98°C for 10 seconds 4) 63°C 30 seconds, repeat steps 3-4 for 20 total cycles, and 5) 72°C for 1 min, and the number of additional PCR cycles needed for each sample, by determining the number of cycles needed to reach 1/3 of the max R. Samples were then amplified for 8-9 total cycles as determined by qPCR. Amplified libraries were run on a 1.5% agarose gel and DNA fragments 150-500 base pairs were extracted and gel purified, and the final library concentration was determined by Qubit High Sensitivity dsDNA quantification kit. ATAC-seq libraries were paired-end sequenced on the Illumina NextSeq500 platform with standard protocols at the UPMC Children’s Hospital of Pittsburgh to a depth of ∼30,000,000 uniquely mapped reads per sample.

## QUANTIFICATION AND STATISTICAL ANALYSES

### TT-seq and RNA-seq data analysis

#### Feature definitions

Protein coding RNAs were defined from protein coding genes using the mm10 Gencode genome annotation V23 ^50^, for a final dataset of 21,596 protein coding genes. uaRNA were defined as the antisense region −1500 base pairs upstream to +500 base pairs downstream of all protein coding transcript TSSs from protein coding genes in the mm10 GENCODE genome annotation. uaRNA regions overlapping with protein coding and long non-coding RNA genes and less than 5 kb downstream of any protein coding gene or annotated long non-coding RNA were removed, and overlapping features were merged for a final dataset of 26,605 regions. Protein coding RNAs and uaRNAs were counted in a strand-specific manner. Enhancer regions were defined with previously described gene-distal DNase 1 hypersensitive sites (GSM1014154^136^), after removing all features within 1 kb of the TSSs annotated mm10 coding genes and those overlapping uaRNA regions. To each remaining DHS, 500 base pairs upstream and downstream were added and merging overlapping regions for a final dataset of 101,587 DHS regions. Enhancers overlapping protein coding genes were counted in a strand-specific manner on the antisense strand, while enhancers within intergenic were counted in a unstranded manner.

Paired-end fastq files were trimmed and adapters were removed with trimmomatic^113^ and cutadapt^114^. Reads were aligned to the mm10 genome using STAR^115^ with options -- outSAMtype SAM, --outFilterMismatchNoverReadLmax 0.02, and --outFilterMultimapNmax 1, and quality filtered using SAMtools^137^ view with options -q 7, -f 2, -bS. Counts were generated using featurecounts^117^, with options -B -t “exon” -g “gene_name” -F GTF -p -s 2 for Gencode features, with options -B -F SAF -p -s 2 for non-coding features, using features as defined. Counts analyses were performed in the R/Bioconductor environment. Raw count matrices were corrected for batch effects using Combat-seq^118^ as indicated in Table S6 with RNA-seq counts adjusted using all Gencode features, and TT-seq counts using all Gencode features and non-coding RNA features while using the experimental condition as the covariate. Differential gene expression analysis was performed using DEseq2^119^ for all available replicates of each condition (n=2 for experimental samples and n=17 for control samples), only keeping features with at least 10 counts in 50% of samples and with option lfcShrink type=”apeglm”. Differentially expressed mRNAs were defined as |log2(FC)|≥0.75 and FDR≤0.05 and differentially transcribed ncRNAs were defined as |log2(FC)|≥0.5 and FDR≤0.05. Gene Ontology analysis on gene sets of interest was performed the R package “clusterProfiler”^120^ using all expressed mRNAs (TPM>0, TT-seq) as background. Strand specific bigwigs were generated with deepTools^121^ using DEseq2-derived sizeFactors with binsize of 1, and averaged bigwigs were generated using all replicates for each condition. Differential bigwigs were generated in deepTools from averaged bigwigs. Browser track images were generated using IGV^122^.

#### mRNA and uaRNA relationship analyses

For each condition, promoters showing significant mRNA and/or uaRNA transcriptional changes were sorted into one of four categories based on transcriptional changes: Only mRNA changes (|log_2_(mRNA change)|≥0.75, |log_2_(uaRNA change)|<0.5), Only uaRNA changes (|log_2_(mRNA change)|<0.75, |log_2_(uaRNA change)|≥0.5), Same directional changes (log_2_(mRNA change)|>=0.75, |log_2_(uaRNA change)|≥0.5 in same direction), or Opposite directional changes (|log_2_(mRNA change)|>=0.75, |log_2_(uaRNA change)|≥0.5 in opposite direction). For promoters in each category, directionality was calculated as the log_10_(mRNA counts/uaRNA counts).

Statistical enrichment for promoters in each category was determined using as *P*≤0.05 with Chi-squared test for homogeneity assuming equal enrichment across all categories.

#### Predicted enhancer promoter pairing analysis

Enhancer promoter pairs (EPPs) were predicted in each condition by defining significantly changed promoters (|log_2_(FC)≥0.75 and FDR≤0.05) and putative enhancers (|log_2_(FC)≥0.5 and FDR≤0.05) changed in the same direction occurring within 100 kb, 250 kb, 500 kb, or 1Mb each other on the same chromosome using bedtools^123^.

We used an approach based on Heger et al (2013)^138^ to generate a distribution of expected EPPs per condition. For each condition, we used a custom script to perform 1000 permutations of randomly shuffling the start and end coordinates of each changed promoter and putative enhancer while preserving the element size and chromosome identity to control for differences in chromosome size, and predicted “expected” EPPs across 100 kb, 250 kb, 500 kb, and 1Mb distance thresholds. *P* values for obtaining the observed number of predicted EPPs were calculated as: (sum of (number of tests with predicted EPPs ≥ observed number of EPPs))/number of tests, and were adjusted using the Benjamini & Yekutieli approach. Enrichment was calculated as: the observed number of predicted EPPs/mean(expected number of EPPs).

#### ATAC-seq analysis

Paired end fastq files were processed using the PEPATAC^124^ pipeline. Briefly, reads were trimmed using trimmomatic^113^, and aligned to the mm10 genome using bowtie2^125^ with options --very-sensitive, -X 2000, and SAMtools^137^ was used to remove reads with MAPQ <10 and mitochondrial reads. Picard^127^ was used to remove PCR duplicates. After alignment, we used the quality control metrics of the PEPATAC^124^ pipeline to evaluate the quality of our ATAC-seq samples to ensure high sample quality. Next, SAMtools was used to merge all replicates of the control and experimental conditions, and then the merged files were then subsampled to the same read depth. Reads were corrected for Tn5 cutting bias by adding +4 bases for positive strand and −5 bases for negative strand using alignmentSieve in deepTools with --ATACshift. Next, peak calling on experimental and control samples was performed by MACS2^126^ with – mode BAMPE --nomodel --shift 75. Using bedTools, peak files from the experimental and control samples were then merged, 50 base pairs on each side were added to each feature, and then overlapping features were again merged to generate a consensus peak set for each depletion condition. Peaks within the consensus peak set were further annotated as promoter or enhancer peaks if they occurred with 1 kb of these features. Overlaps with other peak sets were determined using bedtools.

#### Differential chromatin accessibility analysis

All available replicates were size class filtered for NFR size fragments (1-100 bp) using a custom script, and then all subsampled to the sample read depth using samtools. Reads were then corrected for Tn5 cutting bias using alignmentSieve –ATACshift in deepTools. Read counts on consensus peak sets for each condition were generated using bedtools coverage -counts, and count analyses were performed in R/Bioconductor environment. After removing all peaks with less than 10 counts in 50% of samples, differential accessibility analysis was performed using edgeR for all available replicates of each condition with RUVseq correction. Differentially accessible peaks were defined as |log_2_(FC)|≥0.5 and FDR≤0.05.

#### Tobias analysis

Replicate bam files were merged and subsampled using SAMtools to bring experimental and control merged bam files to the same read depth, and transcription factor foot printing analysis was performed using TOBIAS^81^. Transcription factor motifs were downloaded from the JASPAR^139^ database, and TOBIAS ATACorrect and Score-BigWig were used to generate scored bigwig files across the consensus ATAC-seq peak dataset. TOBIAS BINDetect was used to determine differential binding scores for motifs in experimental and control conditions at promoters and putative enhancers. Only motifs of TFs with TPM≥1 (RNA-seq) and ≥10 binding sites and *P* value ≤0.05 in both conditions at bound and unbound loci were considered.

#### CUT&RUN analysis

Paired end fastq files were trimmed to 25 bp and aligned to the mm10 genome using bowtie2 with options -I 10, -X 1000, -N 1, and --very-sensitive. Picard^127^ was used to filter PCR duplicates. SAMtools was used to filter reads with MAPQ <10, and the reads were sorted into size classes according to the target (1-120 bp for all factors, and 150-500 bp for histone variants post-translational modifications). For peak calling, all replicate bams were merged, and MACS2 was used for peak calling against the corresponding IgG file using -f BAMPE and -q 0.001 for factors or –broad –broad-cut-off 0.001 for histone variants and modifications. All experimental and control peaks were merged to create a consensus peak set. Bedtools was used to define overlap between peak sets of interest. deepTools was used to generate bigwigs using RPGC normalization, --effectiveGenomeSize 2407883318, -e, -bs 5, and --smoothLength 20. deepTools was also used to generate differential bigwigs, heatmaps, and metaplots.

#### ChIP-seq analysis

The full list of all publicly available datasets used in this study can be found in Table S5 and the key resources table. Single end fastq files were trimmed using trimmomatic and aligned to the mm10 genome using bowtie2 with options --best --strata, and reads with MAPQ <10 were removed with SAMtools. For peak calling, all replicate bams were merged, and MACS2 was used for peak calling against the correspond input file using -q 0.001 and -f BAMPE for paired end files or -f BAM for single end files.

#### Differential peak enrichment analysis

CUT&RUN read counts on consensus peak sets for each condition were generated using bedtools coverage-counts, and count analyses were performed in R/Bioconductor environment. After removing all peaks with less than 10 counts in 50% of samples, differential accessibility analysis was performed using edgeR for all available replicates of each condition with RUVseq correction. Differentially enriched peaks were defined as |log_2_(FC)|≥0.5 and FDR≤0.05.

